# Circulating mitochondrial bioenergetics as fingerprint of the hepatic one: how to monitor genetic MASLD

**DOI:** 10.1101/2024.05.06.592717

**Authors:** Erika Paolini, Miriam Longo, Marica Meroni, Paola Podini, Marco Maggioni, Angelo Quattrini, Anna Ludovica Fracanzani, Paola Dongiovanni

## Abstract

**Background & Aims:** Metabolic Dysfunction-Associated Steatotic Liver Disease (MASLD) pathogenesis is shaped by genetics and mitochondrial dysfunction. Recently, we demonstrated that the co-presence of patatin-like phospholipase domain-containing 3 (*PNPLA3*), transmembrane 6 superfamily member 2 (*TM6SF2*) and membrane bound o-acyltransferase domain-containing 7 (*MBOAT7*) polymorphisms predisposes to disease progression in MASLD patients and that their deletion contributes to mitochondrial (mt) maladaptation in an *in-vitro* model. In this work we deepened the impact of genes silencing on mitochondrial dynamism. Then we restored TM6SF2 and/or MBOAT7 wild-type (WT) proteins in the *in-vitro* model to evaluate the rescue of organelles’ morphology/function. Finally, we compared hepatic and peripheral mt-bioenrgetics in MASLD patients carrying PNPLA3, MBOAT7 and/or TM6SF2 loss-of-function variations.

**Methods:** WT proteins were overexpressed through lentiviral transfection, mt-respiration was assessed by Seahorse.

**Results:** The restore of MBOAT7 and/or TM6SF2 wild-type proteins resulted in the assembly of *spaghetti*- shaped mitochondria with improved OXPHOS capacity. Mitochondrial activity was assessed in liver biopsies and peripheral blood mononuclear cells of biopsy-proven (n=44;Discovery cohort) and noninvasively assessed (n=45;Fibroscan-MASLD cohort) MASLD patients stratified according to the presence of the 3 at-risk variants alongside in unrelated liver disease patients (n=45;Unrelated liver disease cohort). In the Discovery cohort, the hepatic bioenergetic profile fully reflecting the circulating one, was impaired in carriers of the risk variants, more so when in combination. We confirmed the lowered serum respirometry in the Fibroscan-MASLD cohort. Finally, the circulating respiration did not change in unrelated liver disease patients, thus demonstrating that it was specifically impaired in MASLD.

**Conclusions:** These results boosted the relevance of mitochondrial circulating respirometry to outline genetically-based MASLD.

## Introduction

Over the past two decades, Metabolic Dysfunction-Associated Steatotic Liver Disease (MASLD) has represented a growing burden on health care worldwide, affecting both children and adults(1, 2). MASLD covers a wide spectrum of chronic liver disorders, ranging from simple steatosis to metabolic dysfunction-associated steatohepatitis (MASH) and fibrosis. In 10–20% of cases, MASH can progress to cirrhosis and hepatocellular carcinoma (HCC), and nowadays, it represents the third leading cause of liver transplantation (LT)(3–5). MASLD is a complex disease whose pathogenesis is shaped by both environmental and genetic factors. Its epidemiology is closely interconnected with epidemic obesity and type 2 diabetes mellitus (T2DM), and it is related to the metabolic syndrome (MetS) which includes an umbrella of metabolic conditions as insulin resistance (IR), dyslipidemia, hyperglycemia, hypertension, and cardiovascular disease (CVD)(6–8).

In the earliest MASLD stages, mitochondria conquer a well-defined role by adopting a strategy called “mitochondrial flexibility” through which they adjust number, mass, and activity by harmonizing their lifecycle and turnover which includes fusion, fission and mitophagy(9, 10). The liver is enriched in mitochondria, ranging from 500 to 4000 number *per* hepatocyte, and these organelles are crucial for the whole-body homeostasis and liver physiology since they provide energy by oxidative phosphorylation (OXPHOS), β-oxidation, tricarboxylic acid (TCA) cycle and ketogenesis(11–14). Mitochondria cannot be generated *de novo* and their shaping leads to the fusion-fission interchange regulated by the peroxisome proliferator-activated receptor (PPAR)-γ coactivators 1 (PGC1α) which also orchestrates OXPHOS capacity, heme biosynthesis, glucose, and lipid metabolism(9, 15). During MASH, the mitochondrial adaptability is lost paralleled by the assembly of misshapen and unfunctional mitochondria thus resulting in impaired β-oxidation, bioenergetic activity, ketogenesis and adenosine triphosphate (ATP) production(9). Additionally, lower levels of mitophagy prompt the hepatic accumulation of failed mitochondria that consequently heighten hepatocellular injury by triggering oxidative stress, inflammation, and release of mitochondrial damage-associated molecular patterns (Mito-DAMPs) among which the mitochondrial DNA (mtDNA) fragments (ccf-mtDNA) are the main component(16).

Recently, an oral, liver-directed, thyroid hormone receptor agonist named Resmetirom has been approved by FDA for the treatment of progressive MASLD. Indeed, data from a Phase 3 trial showed that 29% and 25% of patients who received Resmetirom, had resolution of MASH and improvement of at least one stage of fibrosis, respectively(17), although follow-up studies are required to confirm its efficacy. Resmetirom represents the first and only drug which has been approved for the treatment of MASH. Lifestyle interventions encompassing low caloric diet and physical activity are the still the mainstays of disease treatment, albeit the scant compliance of patients outlines the urgent need for MASLD pharmacotherapy (18). To date, hepatic biopsy is considered the gold standard procedure for the diagnosis and staging of MASLD, since it allows to delineate the degree of steatosis, necroinflammation and ballooning through the scoring system NAFLD Activity Score (NAS)(19, 20). Nonetheless, it exhibits several limitations due to its low applicability, high cost, sample size and repeatability, that in turn could lessen diagnostic accuracy in some patients, including children. Hence, facing the growing prevalence of MASLD alongside the scant diagnostic and therapeutic strategies, newly non-invasive methodological approaches have been proposed to estimate disease severity. Considering the key role of mitochondrial maladaptation in MASLD, emerging evidence highlighted how the release of ccf-mtDNA from hepatocytes into the circulation accurately might estimate organelles’ dysfunction alongside advanced liver injury, thus representing a promising non-invasive tool to identify patients at risk of progressive MASLD(16). In keeping with this theory, the focus in recent years shifted to explore real-time energy changes in both hepatic and peripheral blood(21, 22).

The hereditary component of MASLD has long been established and 50-70% of the individual susceptibility to develop the disease alongside its phenotypic variability are owed to inherited risk factors(23–26). Recently, we have demonstrated that the co-presence of the single nucleotide polymorphisms (SNPs) in patatin-like phospholipase domain-containing 3 (*PNPLA3*), transmembrane 6 superfamily member 2 (*TM6SF2*) and membrane bound o-acyltransferase domain-containing 7 (*MBOAT7*) hugely predisposes to advanced disease, highlighting the relevance of polygenic risk scores to identify at-risk individuals(27–31). The impact of these genetic variations alone or combined over MASLD pathogenesis has been investigated in hepatoma cells (HepG2), homozygous for the I148M PNPLA3 variant which were knocked-out (KO) for *MBOAT7* (MBOAT7^−/−^), *TM6SF2* (TM6SF2^−/−^), or both genes (MBOAT7^−/−^TM6SF2^−/−^) through the clustered regularly interspaced short palindromic repeats/CRISPR-associated protein 9 (CRISPR/Cas9) technology in order to mimic the human proteins loss-of-function. Interestingly, we found that the silencing of these genes and more so when in combination (double KO) disrupts the mitochondrial morphology, increases the organelles content, and triggers the oxidative damage, thus suggesting their involvement in mitochondrial dysfunction which in turn underlines progressive MASLD(27, 32).

Therefore, to deepen the genetics engagement in mitochondrial dysfunction, in the present study we firstly evaluated mitochondrial dynamics in term of fusion/fission balance in HepG2 KO models. Secondly, we overexpressed the wild-type (WT) proteins in KO cells to explore whether TM6SF2 and/or MBOAT7 restoration improves mitochondrial dysfunction in term of morphology and function. To this purpose, we explored the mitochondrial lifecycle and activity alongside hepatocellular damage and metabolic reprogramming. Finally, in the translational perspective to use circulating mitochondrial biomarkers for non-invasively predicting disease severity in genetically predisposed individuals, we compared the hepatic and peripheral bioenergetic profiles in MASLD patients carrying *PNPLA3*, *MBOAT7* and/or *TM6SF2* variations.

## Methods

### Discovery cohort

We evaluated the ROS and H_2_O_2_ production, the mitochondrial complex I, III, IV and citrate synthase kinetic activities as well as respiration capacity in both frozen liver biopsies and peripheral blood mononuclear cells (PBMCs) of 44 MASLD patients who were enrolled consecutively at the Metabolic Liver Diseases outpatient service at Fondazione IRCCS Cà Granda, Ospedale Maggiore Policlinico Milano (Milan, Italy). Inclusion criteria were the availability of a liver biopsy specimen for suspected MASH or severe obesity, DNA samples, and clinical data. MASH was defined by the concomitant presence of steatosis, lobular inflammation, and hepatocellular ballooning. Individuals with excessive alcohol intake (men, >30 g/d; women, >20 g/d), viral and autoimmune hepatitis, or other causes of liver disease were excluded. Informed written consent was obtained from each patient and the study protocol was approved by the Ethical Committee of the Fondazione and conforms to the ethical guidelines of the 1975 Declaration of Helsinki. Demographic, anthropometric, and clinical data of these patients are listed in **Table S1**. MASLD patients were stratified according to the number of risk variants (NRV), as follows: 0 for patients who had no risk variants; PNPLA3, MBOAT7 or TM6SF2 for the presence of GG, TT and TT alleles in homozygous in *PNPLA3*, *MBOAT7* and *TM6SF2* genes, respectively (1 NRV); 3NRV for subjects carrying all 3 at-risk variants either heterozygous or homozygous. PBMCs were extracted from whole blood of MASLD patients and together with liver biopsies were homogenate in MAS buffer as described below.

### Fibroscan-MASLD cohort

We assessed the respiration capacity in frozen PBMCs of 45 patients with non-invasive diagnosis of MASLD performed by ultrasound echography using a convex 3.5 MHz probe and by FibroScan^®^ (Fibroscan-MASLD) at the Metabolic Liver Diseases outpatient service at Fondazione IRCCS Cà Granda, Ospedale Maggiore Policlinico Milano (Milan, Italy). Informed written consent was obtained from each patient and the study protocol was approved by the Ethical Committee of the Fondazione IRCCS Ca’ Granda, Milan and conforms to the ethical guidelines of the 1975 Declaration of Helsinki. Demographic, anthropometric, and clinical data of these patients are listed in **Table S2**. Clinical features of Fibroscan-MASLD cohort matched for age, sex and BMI with patients of the Discovery cohort. Patients were stratified according to the number of risk variant (NRV), as for the Discovery cohort.

### Unrelated Liver Disease Cohort

We evaluated the respiration capacity in frozen PBMCs of 44 patients enrolled at the Metabolic Liver Diseases outpatient service at Fondazione IRCCS Cà Granda (Milan, Italy) and affected by alpha1-antitrypsin deficiency (AAT; n=16), hereditary hemochromatosis (HH; n=16), alcoholic liver disease (ALD; n=7), autoimmune hepatitis (AH; n=5) (MASLD-unrelated cohort). Diagnostic inclusion criteria were the presence of PiZ allele homozygous or heterozygous for AAT, the homozygosity for the C282Y mutation in Homeostatic Iron Regulator (HFE) gene for HH, alcohol intake (>30 g/day for males, >20 g/day for females) for ALD and EASL guidelines for AH(33). Hepatic histology was assessed non-invasively by Fibroscan to estimate the presence of liver steatosis (CAP≥ 248) and fibrosis (LSM ≥ 7.0 and ≥ 6.2 kPa)(34, 35). Informed written consent was obtained from each patient and the study protocol was approved by the Ethical Committee of the Fondazione and conforms to the ethical guidelines of the 1975 Declaration of Helsinki. Demographic, anthropometric, and clinical data of these patients are listed in **Table S3**. Patients belonging to the MASLD-unrelated cohort overlapped clinical features (sex, age and BMI) with MASLD individuals (Discovery cohort, Fibroscan- MASLD cohort) but differentiated in liver disease etiology.

### Statistical Analysis

Statistical analyses were performed using JMP16.0 (SAS, Cary, NC, USA), R statistical analysis version 3.3.2 and Prism (V.9, San Diego, CA, USA), using one-way analysis of variance (ANOVA) or chi-square test where appropriate. For descriptive statistics, continuous variables are shown as mean and standard deviation or median and interquartile range for highly skewed biological variables (i.e. ALT). Variables with skewed distributions were logarithmically transformed before analyses. Categorical variables are presented as number and proportion. Analyses were performed by fitting data to generalized linear regression models. Generalized linear models were fitted to examine continuous traits. Multinomial logistic regression models were fitted to examine binary traits and ordinal regression models were fitted for ordinal traits (components of the MASLD activity score: severity of steatosis, necroinflammation and hepatocellular ballooning, stage of fibrosis). When specified, confounding factors were included in the model. For gene expression analyses, differences between groups were calculated by one-way ANOVA, which was followed by post hoc t-tests adjusted for the number of comparisons when multiple groups were involved (Bonferroni correction). p-values<0.05 (two-tailed) were considered statistically significant.

## Results

### Restoring MBOAT7 and/or TM6SF2 WT activities in KO models rebalances the mitobiogenesis

It is well established that the impaired mitochondrial number and architecture result from unbalanced mitobiogenesis and flow towards maladaptive organelles activity. We recently demonstrated that *PNPLA3*, *MBOAT7* and *TM6SF2* loss-of-functions affects the mitochondrial morphology and biomass in hepatocytes, although we didn’t provide any data regarding their impact on organelle’s dynamics(27). Therefore, in this study we firstly investigated the mechanisms through which the deletion of these genes may impair mitochondrial lifecycle by exploiting HepG2 MBOAT7^−/−^, TM6SF2^−/−^ and MBOAT7^−/−^TM6SF2^−/−^ *in vitro* models.

Then, we restored the WT proteins through a lentiviral transfection in KO cells aiming to obtain stable cell lines that overexpress MBOAT7 (MBOAT7^+/+^) and/or TM6SF2 (TM6SF2^+/+^; MBOAT7^+/+^ TM6SF2^+/+^) genes (**Figure S1A-F**) and to assess whether mitochondrial dynamics, maladaptive function, metabolic reprogramming, and advanced injury may be reversed.

We previously demonstrated that the silencing of MBOAT7 and TM6SF2 genes in HepG2 cells (homozygous for the PNPLA3 I148M mutation) triggered fat accumulation. As evidence that the activity of MBOAT7 and TM6SF2 had been restored, lipid overload, evaluated by ORO staining (**Figure S1G-H)** and triglycerides (**Figure S1I)**/cholesterol (**Figure S1J)** content, was reduced in MBOAT7^+/+^, TM6SF2^+/+^ and MBOAT7^+/+^TM6SF2^+/+^ cells (**Figure S1G-J).**

Next, we went on in evaluating mitochondrial dynamism in KO and overexpressed cells. Protein (**Figure 1A-B**) and mRNA (**Figure S2A**) levels of PGC1α, key regulator of mitochondrial lifecycle and β-oxidation, increased in MBOAT7^−/−^ cells and much more in TM6SF2^−/−^ and MBOAT7^−/−^TM6SF2^−/−^ones (adjusted **(adj-p)<0.01, *adj-p<0.05 *vs* Cas9) as a counter strategy in response to lipid overload (**Figure S1G-J**). In keeping with reduced lipid accumulation, the replenishment of MBOAT7 and TM6SF2 WT proteins declined PGC1α levels showing the strongest effect in MBOAT7^+/+^TM6SF2^+/+^ cell line (**Figure 1A-B, Figure S2A**: **adj-p<0.01 *vs* MBOAT7^−/−^, TM6SF2^−/−^; *adj-p<0.05 *vs* MBOAT7^−/−^TM6SF2^−/−^). Concerning mitochondrial fusion, Mfn1 (**Figure 1A; Figure S2B-F**), Mfn2 (**Figure S2C-G**) and Opa1 (**Figure 1A; Figure S2H**) levels were lower in KO cells and mostly in MBOAT7^−/−^TM6SF2^−/−^ model whereas they were rescued in WT cells and especially in MBOAT7^+/+^TM6SF2^+/+^ ones (*adj-p<0.05 *vs* MBOAT7^−/−^, TM6SF2^−/−^, MBOAT7^−/−^TM6SF2^−/−^). mRNA expression and protein activity of DRP1 (**Figure 1A; Figure S2D-I**) and Fis1 (**Figure 1A; Figure S2E**), which are involved in mitochondrial fission, were higher in MBOAT7^−/−^TM6SF2^−/−^ cells while they returned under physiological condition in WT overexpressed models (*adj-p<0.05 *vs* Cas9, MBOAT7^−/−^TM6SF2^−/−^; **adj- p<0.01 *vs* MBOAT7^−/−^, TM6SF2^−/−^, *MBOAT*7^−/−^TM6SF2^−/−^). Finally, the silencing of MBOAT7 and TM6SF2 alone, and much more in combination, repressed the mitophagy pathway exhibiting vacuolar autophagic structures, that were recovered in WT cells and especially in MBOAT7^+/+^TM6SF2^+/+^ clones (**Figure 1C-F).** Taken together, these data suggest that the deletion of both MBOAT7 and TM6SF2 triggers an imbalance of mitochondrial lifecycle towards high fission and non-operative mitophagy, which explains the increased mass of misfolded organelles in the KO models, also confirmed by confocal microscopy (**Figure 2A-B**). Moreover, we observed that the D-loop copies, reflecting mitochondrial mass, were increased in the silenced models (**Figure 2C**). Conversely, the total number of organelles as well as D-loop-copies was lower in overexpressed models and especially in MBOAT7^+/+^TM6SF2^+/+^ clones (**adj-p<0.01 *vs* Cas9, MBOAT7^−/−^, TM6SF2^−/−^, MBOAT7^−/−^TM6SF2^−/−^) (**Figure 2A-C)** thus suggesting that the recovery of WT proteins resets mitobiogenesis possibly ensuring the assembly of physiological mitochondria (referred to as *spaghetti*-like).

**Figure 1.**
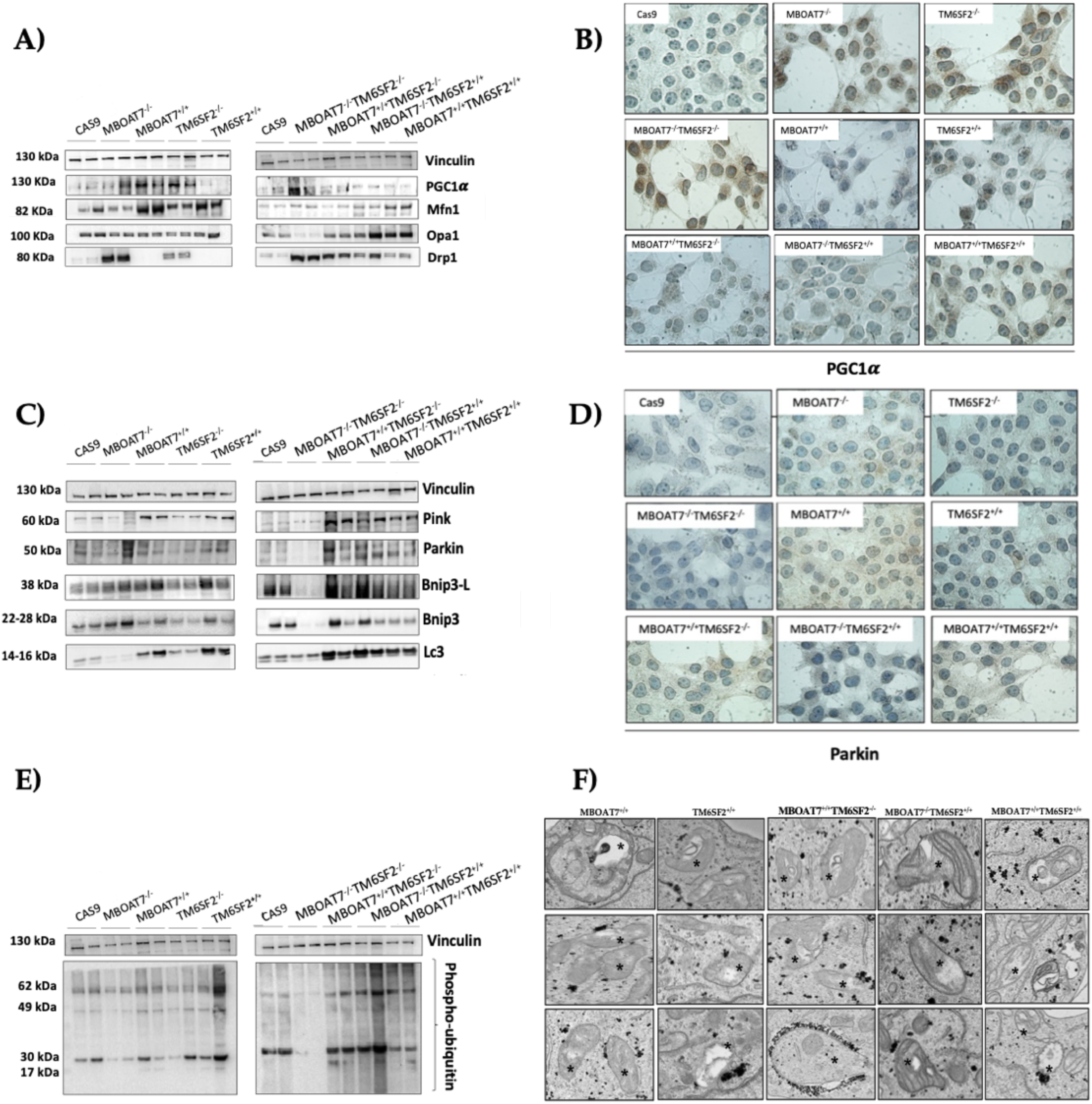
***The MBOAT7 and/or TM6SF2 WT overexpression rebalances the mitochondrial lifecycle and turnover.* A-C-E)** The protein levels of PGC1α, Mfn1, Opa1, Drp1, Pink, Parkin, Bnip3-L, Bnip3, LC3 and phopsho-ubiquitin were assessed by Western blot and normalized to the vinculin housekeeping gene. **B-D)** Cytoplasmatic and nuclear localization of PGC1α and Parkin were represented by Immunocytochemistry pictures. **F)** Representative TEM images of autophagic structures in the form of vacuoles obtained by ultrathin 70-nm sections of hepatocytes. Black stars indicate the autophagic structures in the form of vacuoles in mitochondria and cytoplasm. At least 3 independent experiments were conducted. For bar graphs, data are expressed as means and SD. Adjusted ∗P < .05 and ∗∗P < .01.

**Figure 2.**
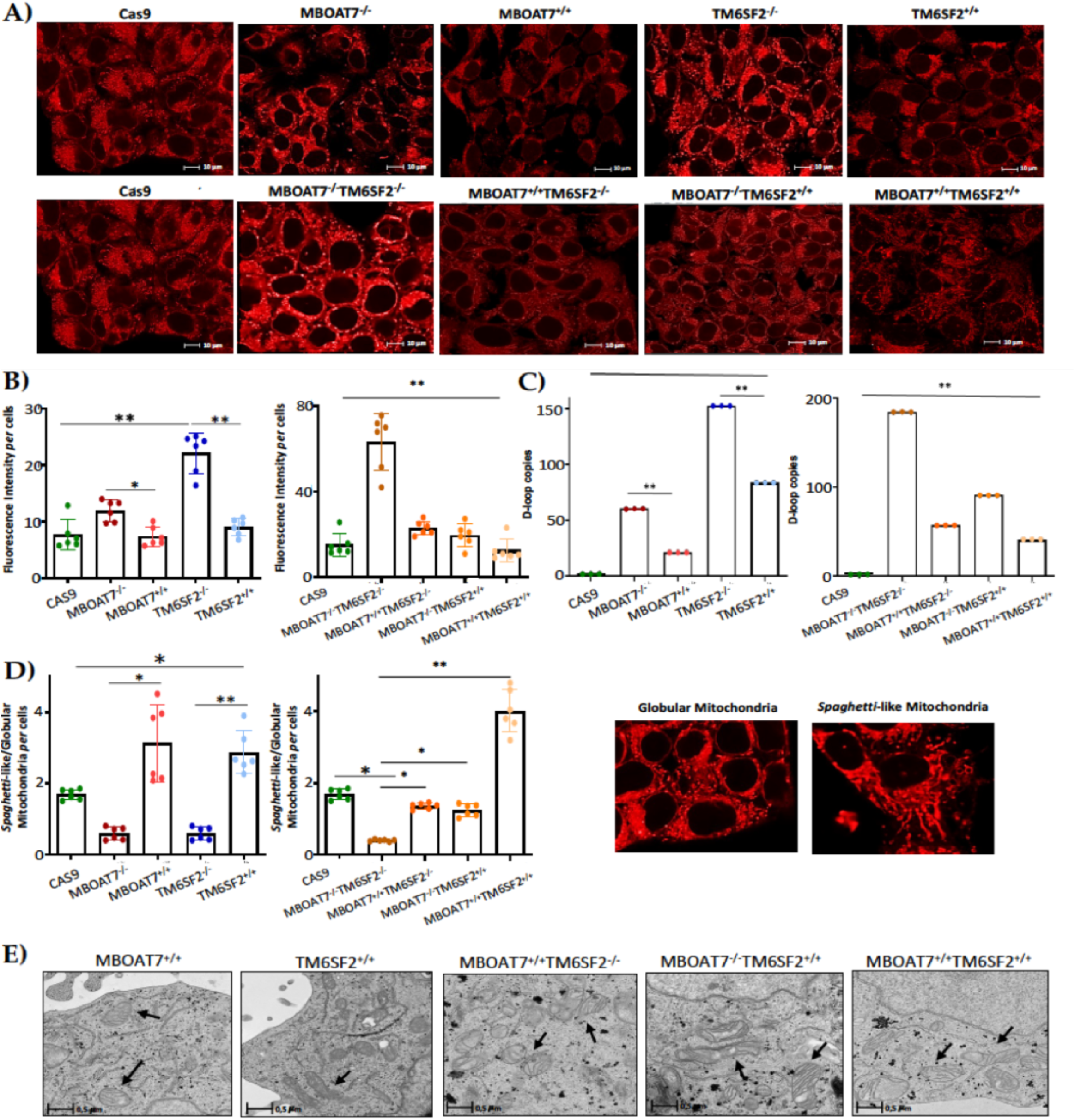
*The WT upregulation of MBOAT7 and/or TM6SF2 WT genes in knock-out models resumes the assembly of physiological shaped mitochondria*. **A)** Representative confocal microscopy images (size range: 10 μm) of mitochondria obtained by Mito Tracker staining. **B)** Fluorescence Intensity per cells was quantified by ImageJ in 6 random nonoverlapping micrographs per condition by calculating the percentage of pixels above the threshold value in respect to total pixels per area. **C)** D-loop copies, that disclose intracellular number of mitochondria, were calculated by reverse-transcription quantitative PCR, and normalized to the RnaseP housekeeping gene. **D)** The ratio between spaghetti-like and globular shaped mitochondria per cells was quantified by ImageJ in 6 random nonoverlapping micrographs per condition by calculating the percentage of pixels above the threshold values of spaghetti-like and globular, respectively, in respect to total pixels per area. **E)** Representative TEM images of mitochondria obtained by ultrathin 70-nm sections of hepatocytes. Black arrows indicate the mitochondria. At least 3 independent experiments were conducted. For bar graphs, data are expressed as means and SD. Adjusted ∗P < .05 and ∗∗P < .01.

### Mitochondrial morphology is restored by the expression of MBOAT7 and/or TM6SF2 wild-type proteins in knock-out models

The aberrant bulk of mitochondria is paralleled by an altered morphology. Indeed, we pointed out that KO cells exhibited many misshapen globular mitochondria compared to the physiological *spaghetti*-like observed in WT overexpressed models (**Figure 2D**), supporting that TM6SF2 and MBOAT7 deletion led to deranged mitobiogenesis paralleled by a suppressed mitophagy (**Figure 1-Figure S2**) (*adj-p<0.05 *vs* Cas9, MBOAT7^−/−^, MBOAT7^−/−^TM6SF2^−/−^; **adj-p<0.01 *vs* TM6SF2^−/−^, MBOAT7^−/−^TM6SF2^−/−^). Consistently with the re-established mitochondrial plasticity in WT overexpressed cells, the organelles appeared more elongated showing regular double membranes and cristae architecture (**Figure 2E**), thus reinforcing the hypothesis according to which the higher number of mitochondria may be related to an imbalance in the life cycle rather than to an enhanced organelles’ function.

### The overexpression of MBOAT7 and/or TM6SF2 WT proteins in KO cells rescues mitochondrial function

To assess whether the overexpression of MBOAT7 and TM6SF2 rescues organelles’ function, we firstly evaluated mRNA levels of *PPAR*α (**Figure 3A**), master regulator of FFAs oxidation. The overexpressed cell lines showed higher expression of *PPAR*α, thus demonstrating that the steatotic phenotype was attenuated after MBOAT7 and/or TM6SF2 rescue (**Figure S1**). Concerning mitochondrial respiration and Krebs cycle, the OXPHOS capacity (**Figure 3B**) alongside the activity of complexes III and IV, ATP synthase and citrate synthase (**Figure S3A- D**) were decreased in KO cells and mainly in MBOAT7^−/−^TM6SF2^−/−^ ones, in keeping with low mitochondrial fusion (**Figure 1A-Figure S2B-C-F-G-H**). Conversely, their levels were restored in WT overexpressed models especially in MBOAT7^+/+^TM6SF2^+/+^ clone in agreement with a balanced mitobiogenesis (**Figure 1-S2**) and physiological mitochondrial plasticity (**Figure 2**) (**adj-p<0.01 *vs* Cas9, MBOAT7^−/−^TM6SF2^−/−^;*adj-p<0.05 *vs* Cas9, TM6SF2^−/−^, MBOAT7^−/−^TM6SF2^−/−^). The quantitative measurement of total ATP rate derived from mitochondrial and glycolytic pathways revealed that the mitochondrial oxygen consumption rate (OCR) which was lower in MBOAT7^−/−^TM6SF2^−/−^ clone, increased in WT overexpressed models and mostly in MBOAT7^+/+^TM6SF2^+/+^ cells (**Figure 3C-D**) (*adj-p<0.05 *vs* Cas9, MBOAT7^−/−^, TM6SF2^−/−^, MBOAT7^−/−^TM6SF2^−/−^). Consistently, the ATP production derived equally from glycolytic pathway and mitochondria in all KO cell lines unless for the MBOAT7^−/−^TM6SF2^−/−^ one which showed the highest extracellular acidification rate (ECAR) as energy source from anaerobic glycolysis (**Figure 3E-F**) (**adj-p<0.001, *adj-p<0.05 *vs* own ATP glycolysis). Contrarywise, the overexpression of MBOAT7 and TM6SF2 alone and mainly combined, enhanced the ATP production related to OXPHOS capacity, thus recovering mitochondrial functions, and impeding the metabolic reprogramming which is hallmark of tumorigenesis (**Figure S4**).

**Figure 3.**
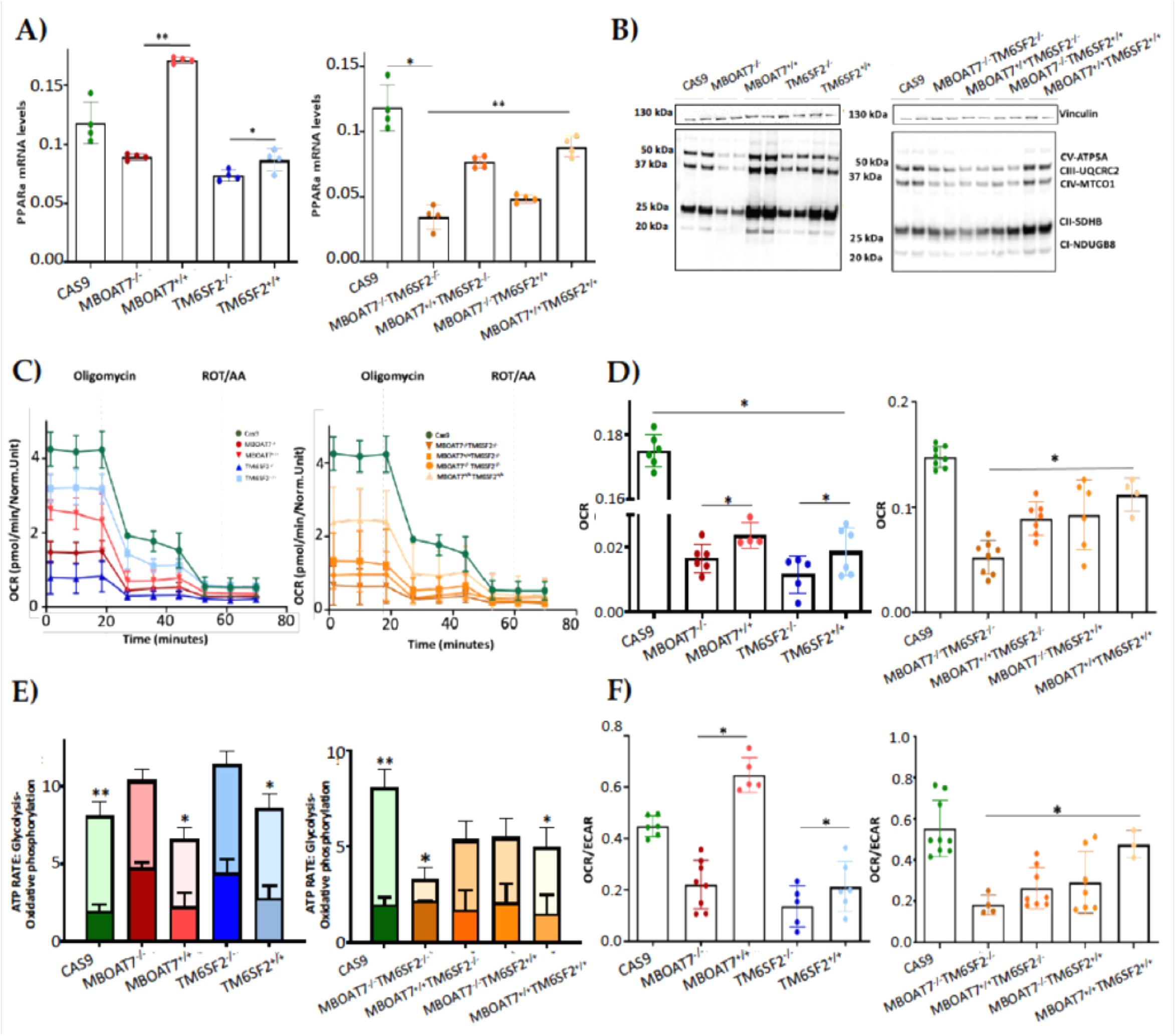
***The MBOAT7 and/or TM6SF2 WT overexpression in knock-out models restores mitochondrial functions.* A)** The mRNA expression of *PPAR*α was evaluated by reverse- transcription quantitative PCR and normalized to the β-actin housekeeping gene. **B)** The protein levels of OXPHOS complexes I, II, III, IV and V were assessed by Western blot and normalized to the vinculin housekeeping gene. **C-D)** The oxygen consumption rate (OCR) was obtained by Mito Stress test using the Seahorse XF Analyzers in live cells. **E-F)** The ATP rate derived from glycolysis and oxidative phosphorylation alongside the OCR/ extracellular acidification rate (ECAR) ratio were measured by Glycolytic Assay exploiting the Seahorse XF Analyzers in live cells. At least 3 independent experiments were conducted. For bar graphs, data are expressed as means and SD. Adjusted ∗*P* < .05 and ∗∗*P* < .01.

### The restore of MBOAT7 and/or TM6SF2 WT proteins in KO cells attenuates hepatocellular damage

The switch toward anaerobic glycolysis (**Figure 3-Figure S3**) is one of the first key step in the malignant transformation paralleled by high proliferation capacity and invasiveness (**Figure S4**). In support to this notion, KO models and particularly the MBOAT7^−/−^TM6SF2^−/−^ clone converted glucose into lactate by triggering lactate dehydrogenase (LDH) activity and production, both reduced in WT overexpressed models (**Figure 4A- C**) (*adj-p<0.05 *vs* Cas9, TM6SF2^−/−^; **adj-p<0.01 *vs* MBOAT7^−/−^, MBOAT7^−/−^TM6SF2^−/−^). Lactate directly takes part in oxidative damage resulting in boosted Reactive Oxygen Species (ROS)/Reactive nitrite species (ROS/RNS) ratio, ROS-induced DNA damage and malondialdehyde (MDA) production which were higher mostly in MBOAT7^−/−^TM6SF2^−/−^ cells, whereas they strongly decreased after the MBOAT7 and/or TM6SF2 WT upregulation (**Figure 4D-F**) (*adj-p<0.05, **adj-p<0.01 *vs* Cas9, MBOAT7^−/−^, TM6SF2^−/−^, MBOAT7^−/−^TM6SF2^−/−^). As a response to unbalanced mitobiogenesis (**Figure 1-S2**) and failed mitochondrial activity (**Figure 3-S3**), KO models released higher circulating cell-free mitochondrial DNA (ccf-mtDNA) which derives from damaged organelles (**Figure 4G**). Otherwise, the upregulation of MBOAT7 and/or TM6SF2 WT proteins greatly reduced the release of ccf-mtDNA (**Figure 4G**), corroborating the hypothesis that the high number of mitochondria is not representative of an enhanced function and that genetics directly impacts on the integrity of mitochondria. Notably, ccf-mtDNA correlated with lower activity of complex V (ATP synthase) (p<.0001, β=-1.1, 95%CI, -1.56^-^-0.64) and reduced OCR (p<.0001, β=-9.9, 95%CI, -16.77^-^-3.10). Hence, the assessment of ccf-mtDNA levels may mirror mitochondrial derangement due to *TM6SF2* and *MBOAT7* deletion, thus paving the way to consider them as a mitochondrial circulating biomarker of progressive MASLD in genetically predisposed individuals.

**Figure 4.**
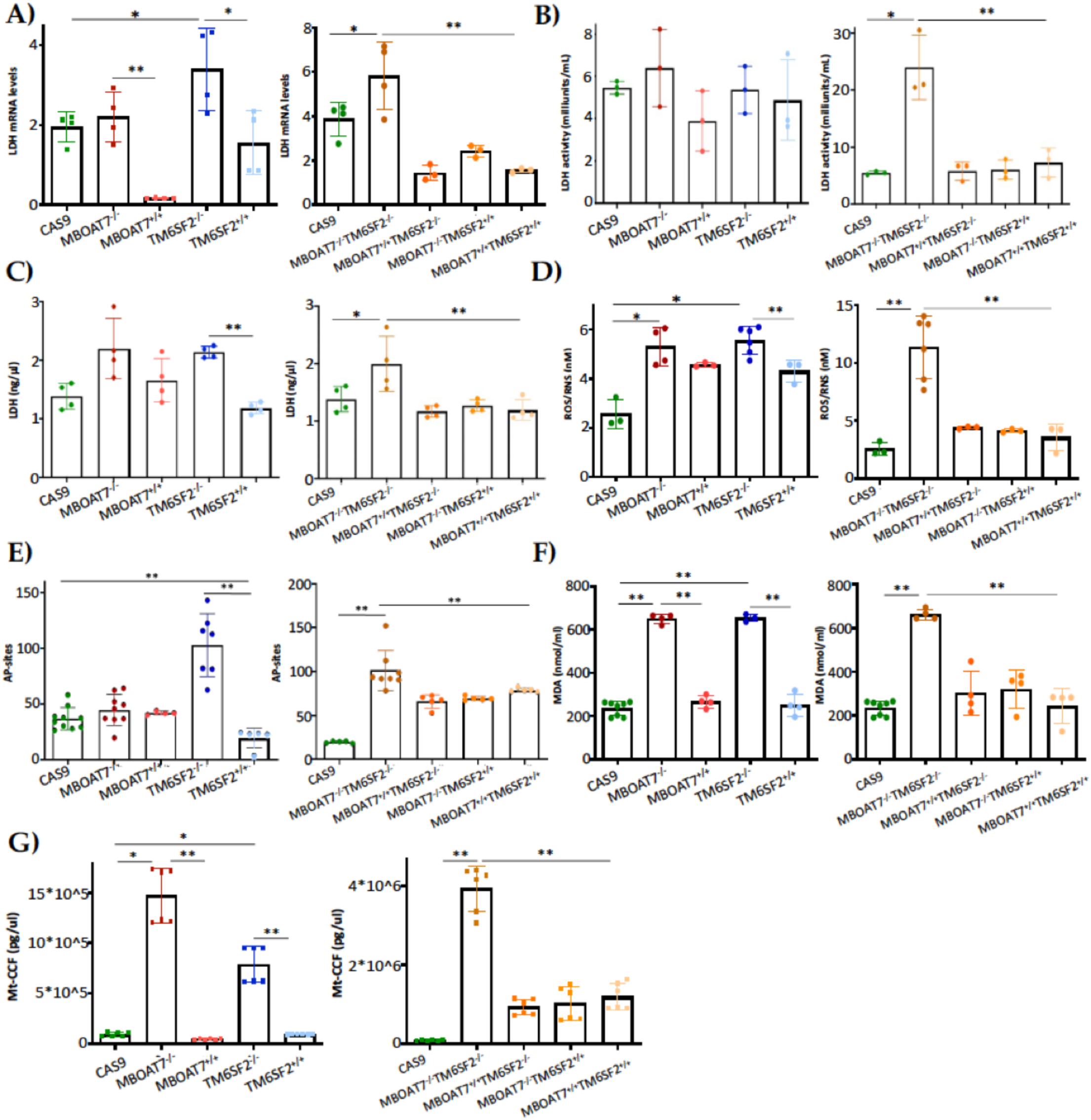
***The MBOAT7 and/or TM6SF2 WT upregulation in knock-out cells attenuates hepatocellular injury.* A)** The mRNA expression of *LDH* was evaluated by reverse-transcription quantitative PCR and normalized to the β-actin housekeeping gene **B)** The LDH activity was measured by the Lactate Dehydrogenase (LDH) Assay Kit in cell lysates. **C)** The LDH production was quantified through the LDH activity by Lactate Dehydrogenase (LDH) Assay Kit in cell supernatants. **D)** The oxidative stress was evaluated by Dichlorodihydrofluorescin (DCF) Reactive oxygen species/Reactive nitrogen species (ROS/RNS) Colorimetric Assay Kit in cell lysates. **E)** The ROS-induced DNA damage was detected through the DNA Damage Colorimetric Assay Kit (apurinic/apyrimidinic (AP) sites) in cell lysates. **F)** The malondialdehyde (MDA) production was calculated by the Lipid Peroxidation (MDA) Assay Kit in cell lysates. **G)** The release of cell- free mitochondrial DNA fragments (mt-ccf) was quantified through quantitative real-time PCR and normalized on the standard curve obtained from serial dilutions of a sample pool at known concentration. At least 3 independent experiments were conducted. For bar graphs, data are expressed as means and SD. Adjusted ∗*P* < .05 and ∗∗P < .01

### The PNPLA3 I148M overexpression in hepatoma cells impairs the mitochondrial function

In our MASLD *in vitro* models, we deeply demonstrated that MBOAT7 and TM6SF2 loss-of-functions in HepG2 cells triggered to mitochondrial maladaptation, exhibiting the strongest effect when both genes were deleted (MBOAT7^-/-^TM6SF2^-/-^). Moreover, the restoration of MBOAT7 and TM6SF2 WT activities rescued the mitochondrial phenotype, corroborating their role in contributing to mitochondrial dysfunction encompassing lifecycle, morphology and activity.

As HepG2 cells carry the PNPLA3-I148M variant in homozygosity, we were not able to discriminate its effect on mitochondrial morphology and function from that of TM6SF2 and MBOAT7 in KO models. Interestingly, Cas9 cells carrying the I148M protein did not affect the organelles dynamics and function, hinting that PNPLA3 loss-of-function mutation may not play a role in the mitochondrial dysfunction. Nonetheless, literature evidence showed that the overexpression of I148M in hepatoma cells was correlated with mitochondrial impairment (36). Therefore, to clarify the impact of PNPLA3 variation on mitochondrial dynamics, we performed a lentiviral transfection in HepG2 (in order to force the mutation effect) and Hep3B (which are WT for the PNPLA3-I148M mutation) cells by using pLenti-C-mGFP-P2A-Puro lentiviral vectors (HepG2 I148M^+/+^and Hep3B I148M^+/+^). The vector design embraced a GFP tag fused with an ORF (PNPLA3-I148M) targeting the I148M protein that was upregulated in all cell lines (**Figure 5A-B**) (**adj-p<0.01 *vs* HepG2 and Hep3B). At ORO staining, lipid accumulation increased in Hep3B I148M^+/+^ cells and more so in HepG2 I148M^+/+^ ones, resulting in the assembly of larger LDs (**Figure 5C**). Concerning mitobiogenesis, PGC1α protein levels augmented in both overexpressed models suggesting a responsive strategy to counter fat accumulation (**Figure 5D**). About fusion, Opa1 activity was reduced in the overexpressed cell lines (**Figure 5D**) supported by lower OXPHOS capacity observed at Western Blot (**Figure 5E**). In keeping with these results, ATP synthase activity, ATP content and citrate synthase activity decreased in all overexpressed cells, emphasizing that the I148M overexpression dampened the oxidative phosphorylation probably due to high lipid overload (**Figure 5F-H**) (***adj-p<0.001, **adj-p<0.01 and *adj-p<0.05 *vs* HepG2 and Hep3B). Consistently, the ROS/RNS ratio (**Figure 5I)** and LDH production (**Figure 5J)** were boosted in cells overexpressing the mutated protein alongside the release of ccf-mtDNA (**Figure 5K)** (***adj-p<0.001, **adj-p<0.01 and *adj-p<0.05 *vs* HepG2 and Hep3B). To sum up, the upregulation of I148M-PNPLA3 protein in *in vitro* hepatoma cells, by resembling the overload of mutated protein in human carriers, unveiled its involvement in mitochondrial dysfunction.

**Fig 5.**
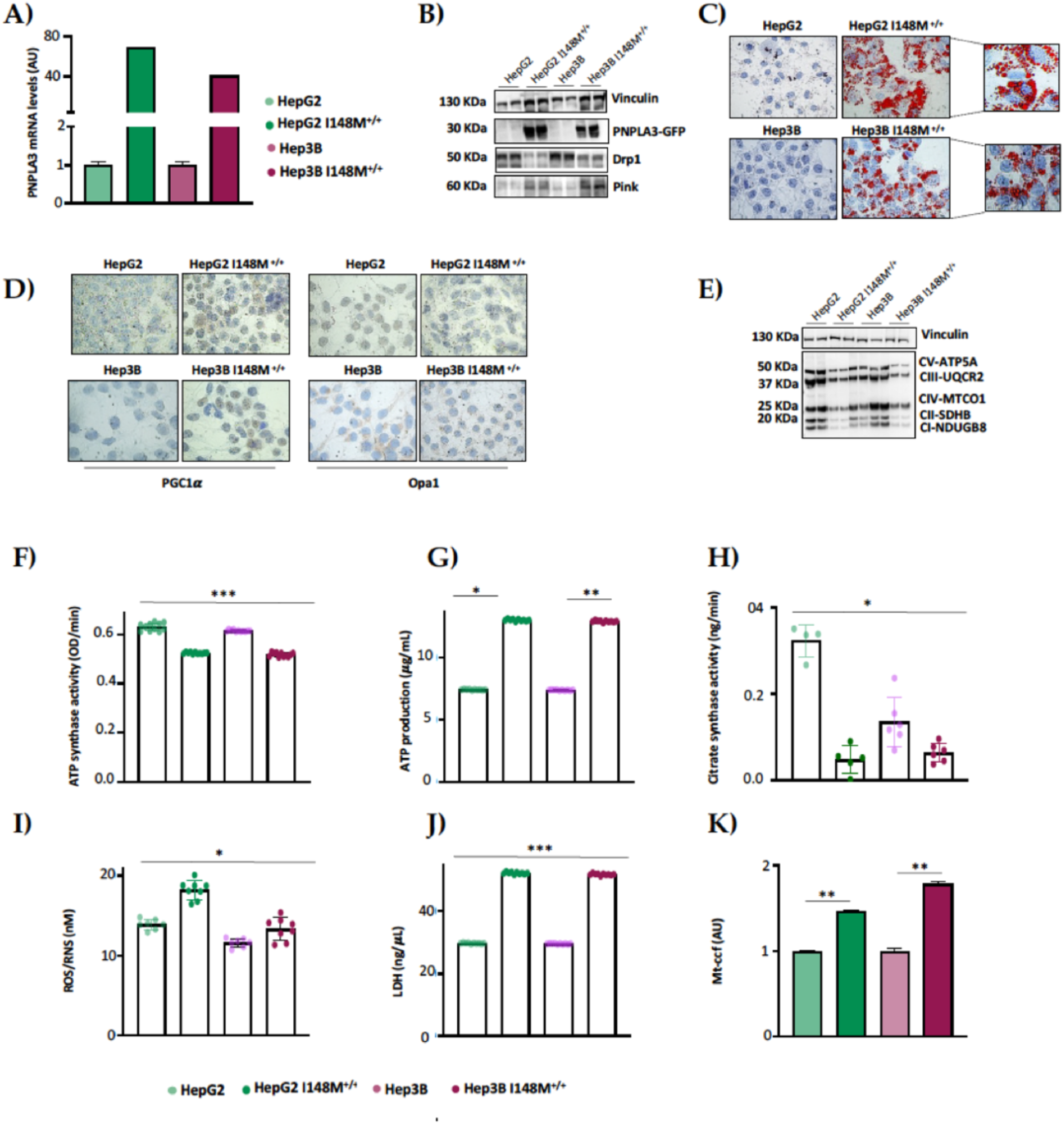
***The I148M PNPLA3 upregulation in hepatoma cells triggers lipid accumulation alongside mitochondrial failure.* A)** The mRNA expression of *PNPLA3* was evaluated by reverse-transcription quantitative PCR and normalized to the β-actin housekeeping gene. **B)** The protein levels of PNPLA3 tagged with GFP (PNPLA3-GFP) were assessed by Western blot and normalized to the vinculin housekeeping gene. **C**) LD accumulation was assessed by ORO staining (magnification, 630×). **D)** Cytoplasmatic and nuclear localization of PGC1α and Opa1 were represented by Immunocytochemistry pictures. **E)** The protein levels of OXPHOS complexes I, II, III, IV and V were assessed by Western blot and normalized to the vinculin housekeeping gene. **F)** The ATP5A (mtDNA–encoded subunit of the complex V) activity was quantified by ATP Synthase Enzyme Activity Microplate Assay Kit in isolated mitochondria from cell lysates. **G)** The ATP production was quantified by ATP Colorimetric Assay in cell lysates. **H)** The citrate synthase activity was assessed through the Citrate Synthase Assay Kit in isolated mitochondria from cell lysates. **I)** The ROS-induced DNA damage was detected through the DNA Damage Colorimetric Assay Kit (apurinic/apyrimidinic (AP) sites) in cell lysates. **J)** The LDH production was quantified through the LDH activity by Lactate Dehydrogenase (LDH) Assay Kit in cell supernatants. **K)** The release of cell-free mitochondrial DNA fragments (mt-ccf) was quantified through quantitative real-time PCR and normalized on the standard curve obtained from serial dilutions of a sample pool at known concentration At least 3 independent experiments were conducted. For bar graphs, data are expressed as means and SD. Adjusted ∗*P* < .05 and ∗∗*P* < .01.

### Hepatic and circulating mitochondrial activity is impaired in MASLD patients carrying the 3 at risk variants

Based on our *in vitro* results, the co-presence of *PNPLA3*, *MBOAT7* and *TM6SF2* loss-of-functions in HepG2 cells impacts on mitochondrial plasticity exhibiting the restore of organelles’ dynamics and function after the overexpression of WT proteins. The mitochondrial impairment is mirrored by the release of ccf-mtDNA. All in all, these genes have been identified as the main predictors of progressive MASLD which is featured by damaged mitochondria which in turn release ccf-mtDNA. Therefore, in order to translate these *in vitro* findings into clinic and to consider the assessment of mitochondrial molecules as non-invasive biomarkers, we evaluated and compared the mitochondrial activity in frozen liver biopsies and peripheral blood mononuclear cells (PBMCs) in 44 MASLD patients (Discovery cohort). Patients were stratified according to genetic background as follows: WT (n=11; 22.7%), homozygous for the I148M PNPLA3 (n=11; 25.0%), homozygous for the rs641738 MBOAT7 (n=9, 20.5%), homozygous for the E167K TM6SF2 (n=7, 15.9%), carrying all 3 at-risk variants (3NRV) in heterozygosis or homozygosis (n=7, 15.9%) (**Table S1**). Hepatic ROS (**Figure 6A-Table S6**) and H_2_O_2_ (**Figure 6B- Table S6**) levels increased in GG *PNPLA3*, TT *MBOAT7* and TT *TM6SF2* carriers and even more in 3NRV ones (*p<0.05, **p<0.001 *vs* 0NRV). Interestingly, a comparable result was observed in PBMCs homogenate suggesting that their mitochondrial performance may reflect the hepatic one (**Figure 6A-B**). Concerning mitochondrial respiration, the total and kinetic activity of citrate synthase (**Figure 6C- Table S6- S7**), mitochondrial complex I (**Figure 6D- Table S6-S7**), mitochondrial complex III (**Figure 6E- Table S6-S7**) and ATP synthase (**Figure 6F- Table S6-S7**) were lower in both liver biopsies and PBMCs of GG *PNPLA3*, TT *MBOAT7* and TT *TM6SF2* individuals showing the main reduction in 3NRV carriers (*p<0.05, **p<0.001, ***p<0.0001 *vs* 0NRV). Therefore, we firstly demonstrated that in genetically susceptible MASLD patients the mitochondrial failure in the PMBC closely mirrors the hepatic one.

**Figure 6.**
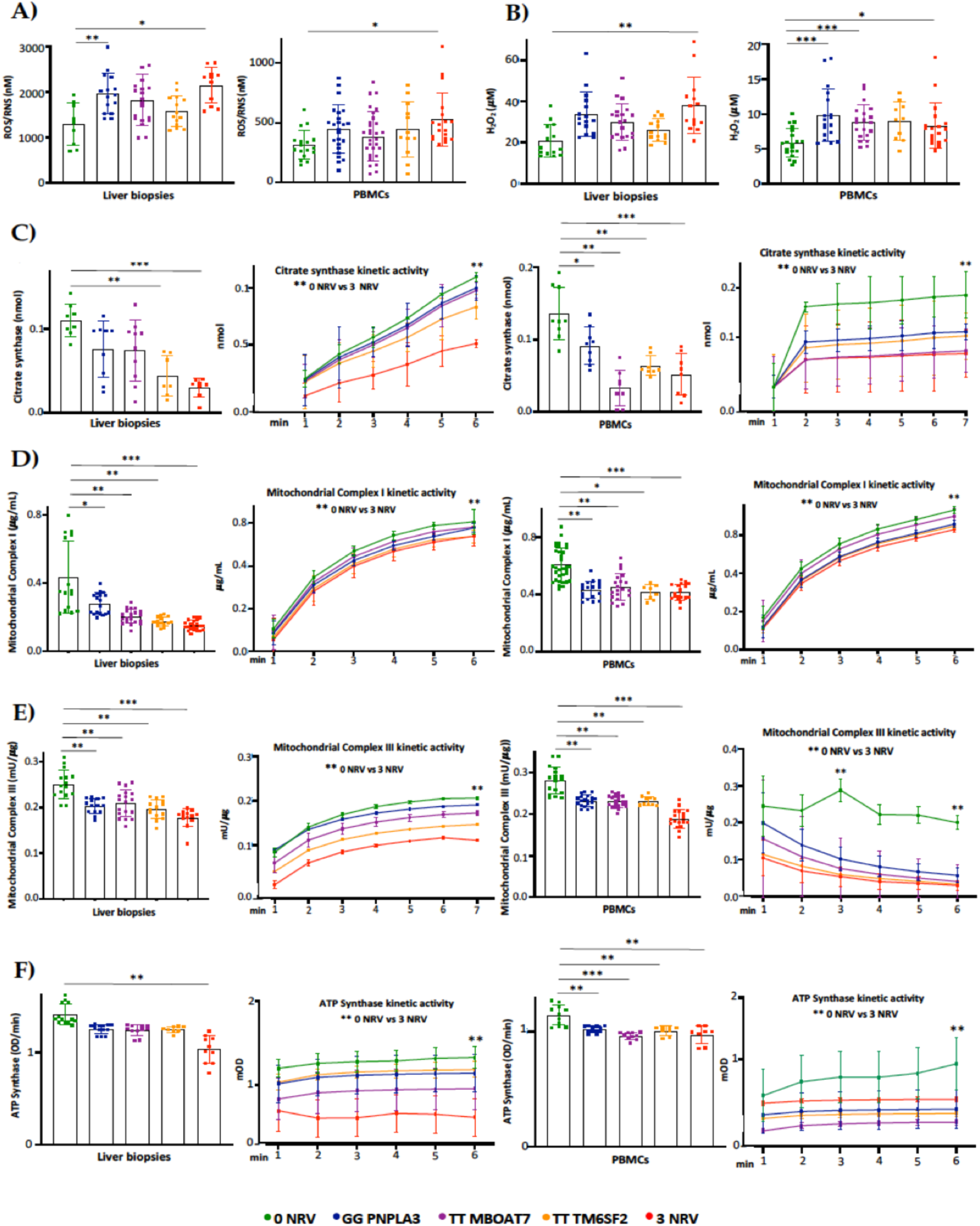
The mitochondrial activity is impaired in liver biopsies and PBMCs of 3NRV MASLD carriers. A-**B)** The oxidative stress was evaluated by Dichlorodihydrofluorescin (DCF) Reactive oxygen species/Reactive nitrogen species (ROS/RNS) Colorimetric Assay Kit in frozen liver biopsies and PBMcs. **C-D)** The complex I/III enzymatic activity was measured through a colorimetric assay in in frozen liver biopsies and PBMcs. **E)** The ATP5A (mtDNA–encoded subunit of the complex V) activity was quantified by ATP Synthase Enzyme Activity Microplate Assay Kit in in frozen liver biopsies and PBMcs. At least 3 independent experiments were conducted. For bar graphs, data are expressed as means and SD. Adjusted ∗*P* < .05 and ∗∗*P* < .01

### Serum bioenergetic profile resembles the hepatic one and prognose MASLD severity in 3NRV carriers

In keeping with the impaired mitochondrial activity, we demonstrated that hepatic and sera OCR progressively decreased from I148M PNPLA3 carriers (30%), to rs641738 MBOAT7 (70%) or E167K TM6SF2 (80%) variations with the lowest levels in 3NRV individuals (90%) (***p<0.0001 *vs* 0NRV) (**Figure 7A-B**), thus resembling the declined OXPHOS capacity observed in MBOAT7^−/−^TM6SF2^−/−^ model (**Figure3-S3**). At multivariable analysis adjusted for age, sex, BMI and diabetes, the co-presence of 3NRV was independently associated with reduced respiration in terms of complex I/IV (**Table S8.** Liver biopsies: p=0.0001, β =-43.37, 95%CI, -64.81^-^-21.92; PBMCs: p=0.002, β=-30.7, 95%CI, -49.96^-^-11.45) and II/IV in both hepatic samples and sera (**Table S9**. Liver biopsies: p=0.0001, β=-27.24, 95%CI, -10.47^-^-14.02 PBMCs: p<0.0001, β =-20.08, 95%CI, -25.20^-^-14.96). These data highlight that genetics directly compromises the circulating bioenergetic profile which closely reflects the hepatic mitochondrial respiration. To investigate whether impaired OCR could estimate mitochondrial failure and thereby progressive disease, patients belonging to the Discovery cohort were stratified according to the presence of MASH-fibrosis. Liver biopsies and PBMCs samples of MASH-Fibrosis patients exhibited a significant OCR reduction in terms of complex I/IV and II/IV activities (**Figure 7C**) (*p, **p at two-way ANOVA). At multinomial logistic regression analysis adjusted for 3NRV, the hepatic respiration capacity of complexes I/IV and II/IV displayed a prognostic value for MASH-Fibrosis of 76% and 79%, respectively (**Figure 7D**). Intriguingly, the sera OCR-I/IV and OCR-II/IV revealed a competitive prediction capacity for MASH-Fibrosis of 85% and 84% (**Figure 7D**). Therefore, we confirmed that circulating mitochondrial respiration reflects that observed in the liver thus making reasonable the applicability of mitochondrial molecules (i.e. ccf-mtDNA) as non-invasive biomarkers of advanced liver disease in patients with a genetic predisposition.

**Figure 7.**
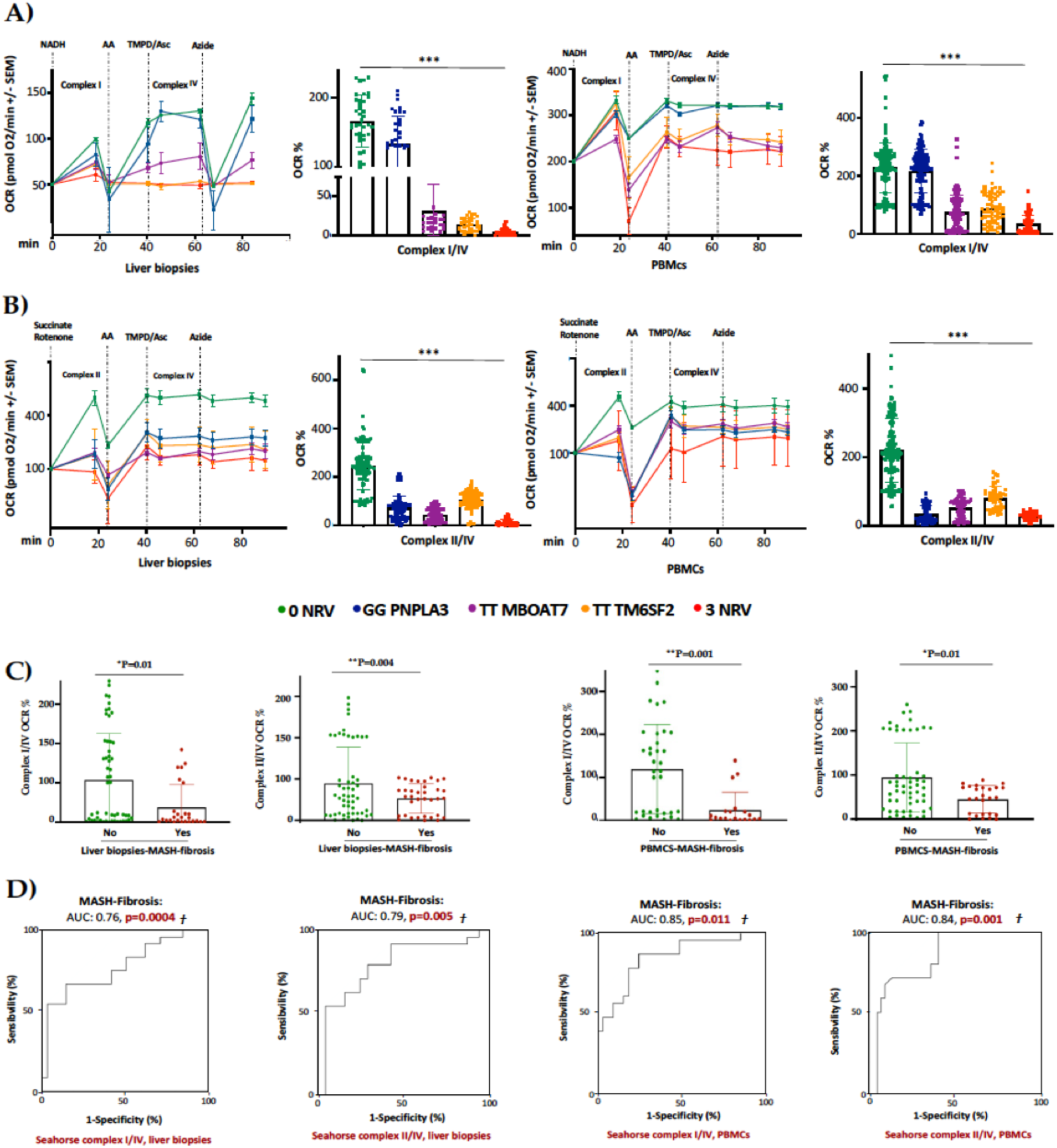
The PBMCs bioenergetic profile reflects the hepatic one and exhibits a strong prognostic value for MASH-Fibrosis in 3NRV carriers. A-B) The oxygen consumption rate (OCR) was obtained by customized Seahorse assay using the Seahorse XF Analyzers in frozen hepatic biopsies and PBMCs. At least 4 independent experiments were conducted. For bar graphs, data are expressed as means and SD. Adjusted ∗*P* < .05 and ∗∗*P* < .01. **C)** At bivariate analysis patients with MASH-fibrosis exhibit lower OCR (Liver biopsies complexI/IV: P=0.01; Liver biopsies complexII/IV: P=0.004; PBMCs complexI/IV: P=0.001; PBMCs complexII/IV: P=0.01). **D)** Multinomial logistic regression analysis adjusted for 3NRV show that complexes I/IV and II/IV have a strong prognostic accuracy for MASH-Fibrosis (Liver biopsies complexI/IV: AUC=0.76, P=0.0004; Liver biopsies complexII/IV: AUC=0.79, P=0.005; PBMCs complexI/IV: AUC=0.85, P=0.011; PBMCs complexII/IV: AUC=0.84, P=0.001)

### Serum bioenergetics predicts fibrosis in non-invasively assessed MASLD patients carrying 3NRV

Next, we evaluated whether the PBMC mitochondrial respiration may predict advanced disease also in patients with a non-invasive diagnosis of MASLD. To this purpose we measured serum OCR in 45 MASLD patients included in the Fibroscan-MASLD cohort. The PBMCs respiration hugely declined in rs641738 MBOAT7 and E167K TM6SF2 carriers and mainly in 3NRV individuals (***p<0.0001 *vs* 0NRV) (**Figure 8A**), thus mirroring the OCR genetic-based signature observed in biopsied patients (Discovery cohort) (**Figure 7B**). At multivariable analysis adjusted for age, sex, BMI and diabetes, the co-presence of 3NRV was independently associated with reduced serum respiration (**Table S10.** p<.0001, β=-63.06, 95%CI, -85.42^-^-40.7), corroborating the direct role of genetics in impairing PBMCs respirometry. To validate that the altered circulating OCR could estimate advanced hepatic disease, these patients were stratified according to the presence of fibrosis (Stiffness >7 kPa). The latter was higher in 3NRV carriers (n=9, 87.50%) than 0NRV (n=0, 0%) and 1NRV (n=6, 23.81%) (**Figure 8B**).

**Figure 8.**
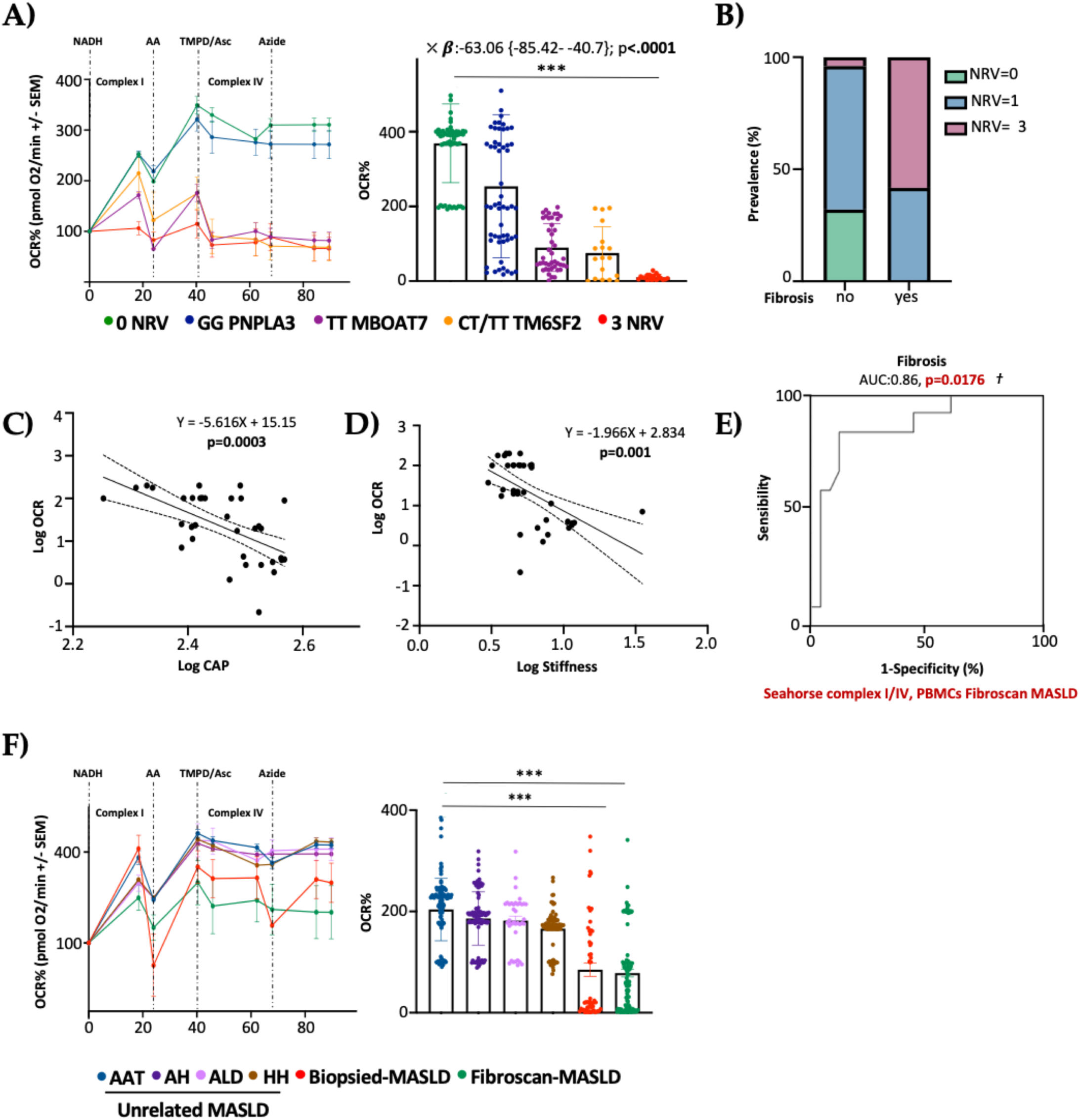
The PBMCs bioenergetic profile predicts fibrosis in both non-invasively and biopsied MASLD patients carrying the at-3 risk variants. A) The oxygen consumption rate (OCR) was obtained by customized Seahorse assay using the Seahorse XF Analyzers in frozen PBMCs. At least 4 independent experiments were conducted. For bar graphs, data are expressed as means and SD. Adjusted ∗*P* < .05 and ∗∗*P* < .01. At multivariable analysis adjusted for age, sex, BMI and diabetes, the co-presence of 3NRV was independently associated with reduced respiration (**X:** p<.0001, β = -63.06, 95%CI, -85.42 to -40.7) **B)** Patients were stratified according to Fibroscan evaluation as fibrosis no= stiffness <7 kPa (n=30) and fibrosis yes= stiffness >7 kPa (n=15). Based on genetic variations and histological assessment, the subjects were grouped as follow: NRV=0, fibrosis, no (n=9, 100%); NRV=1, fibrosis, no (n=20, 76.19%), fibrosis yes (n=6, 23.81%); NRV=3, fibrosis, no (n=1, 12.50%), fibrosis yes (n=9, 87.50%). The prevalence (%) was calculated on the total number of patients/group. **C-D)** Linear correlation analyses of liver steatosis (CAP) and fibrosis (stiffness) scores with oxygen consumption rate (OCR). Overall, the Fibroscan-MASLD patients were included, and the simple linear regression equation and corresponding p value were shown in the figures. **E)** Complexes I/IV have a strong prognostic accuracy for fibrosis (stiffness >7 kPa) (PBMCs complexI/IV: AUC=0.86, P=0.0176) at multinomial logistic regression analysis adjusted for 3NRV. **F)** The oxygen consumption rate (OCR) was obtained by customized Seahorse assay using the Seahorse XF Analyzers in frozen PBMCs. At least 4 independent experiments were conducted. For bar graphs, data are expressed as means and SD. Adjusted ∗*P* < .05 and ∗∗*P* < .01.

Moreover, serum OCR inversely correlated with CAP-steatosis and stiffness-fibrosis scores (**Figure 8C**: Log OCR *vs* Log CAP p=0.0003; **Figure 8D**: Log OCR vs Log stiffness p=0.001), exhibiting a strong prognostic value for fibrosis at ROC curve (86%) (**Figure 8E)** and confirming the results obtained in biopsied patients.

As expected, the presence of 3 at risk variants (3NRV) independently correlated with failed OCR in PBMC when we combined the Discovery and Fibroscan-MASLD cohorts (Overall cohort n=89) (**Table S11.** p<.0001, β=-79.05, 95%CI, -96.29^-^-61.82). Moreover, in the attempt to define the contribute of the individual mutations, we observed a stronger impact of MBOAT7 and TM6SF2 SNPs on circulating respirometry (**Table S11.** MBOAT7 T allele, yes: p<.0001, β=-30.46, 95%CI, -44.64^-^-16.28; TM6SF2 T allele, yes: p=0.0002, β=-29.02,

95%CI, -43.76^-^-14.28). Finally, to deepen whether serum OCR may represent a specific biomarker of MASLD severity (also independently from genetics), we evaluated mitochondrial respiration in PBMCs of 44 patients affected by unrelated liver diseases encompassing AAT, HH, ALD and autoimmune hepatitis (MASLD- unrelated cohort). Serum OCR was similar between the unrelated liver diseases and in all these etiologies OCR was higher compared to that observed in both biopsied-Discovery and Fibroscan-MASLD cohorts (***p<0.0001 *vs* Biopsied-MASLD and Fibroscan-MASLD) (**Figure 8F**). At multivariable analysis adjusted for age and sex, a lower serum OCR was definitively associated with both biopsied and noninvasively assessed MASLD, thereby highlighting the impaired serum respirometry is specific of MASLD (**Table S12.** Biopsied- MASLD *vs* Unrelated MASLD: p=0.01, β=-19.61, 95%CI, -34.56^-^-4.65; Fibroscan-MASLD *vs* Unrelated MASLD: p=0.0008, β=-20.29, 95%CI, -31.82^-^-8.75; Overall MASLD *vs* Unrelated MASLD: p=0.03, β=-16.61, 95%CI, -31.76^-^-1.46).

## Discussion

In the present study, we examined the relationship between genetics and mitochondrial dysfunction, both guilty players in MASLD pathogenesis(9, 23, 27). Nowadays, MASLD is the most common chronic liver disorder worldwide and liver biopsy still remains the gold standard method for the diagnosis of its progressive forms. Facing the growing MASLD prevalence alongside scant diagnostic and therapeutic strategies, newly non-invasive methodological approaches have been proposed to estimate the severity of liver disease(16). One of the key steps which features the MASLD switch towards MASH is represented by mitochondrial dysfunction due to heighten fat accumulation. This results in unbalanced mitochondrial lifecycle that translates in aberrant mitochondrial number, morphology, and activity worsening progressive liver damage. The disequilibrium of mitobiogenesis alongside the consequent accumulation of failed mitochondria have been observed in hepatic tissues of both MASH individuals and murine models, thereby underling that MASLD may be considered as a mitochondrial disorder(9). In addition, it has been described that a high release of ccf-mtDNA from damaged liver organelles along with altered serum respirometry correlates with progressive MASLD, thus suggesting the potential use of circulating mitochondrial molecules as noninvasive biomarkers(16, 22).

MASLD is a multifactorial disease and exhibits a strong hereditable component(23, 37, 38). We demonstrated that the co-presence of *PNPLA3*, *MBOAT7* and *TM6SF2* at-risk variants in 1380 MASLD subjects hugely predisposes to HCC development, highlighting the usefulness of polygenic risk scores to predict disease severity. By exploiting a genetic *in vitro* model, we demonstrated that the deletion of *MBOAT7* and *TM6SF2*, alone and mostly combined, overflowed intracellular lipid accumulation and in parallel triggered mitochondrial derangement in terms of morphology and activity, thereby contributing to advanced hepatic damage(27).

Since MASLD occurs as a mitochondrial disease and PNPLA3, MBOAT7 and TM6SF2 loss-of-functions seem to achieve a role in organelles’ dysfunction, in this study we firstly investigated the impact of genes deletion on mitochondrial dynamism and integrity in KO HepG2 cells. Then we explored whether the restoring of the WT proteins in this *in vitro* model could rescue mitochondrial activity and morphology and dampen the release of molecules such as ccf-mtDNA. Finally, we compared the hepatic bioenergetic profile with the circulating one in patients carrying the three at risk variants with the purpose to define a prognostic and specific signature of progressive MASLD in these individuals.

We observed an impaired mitochondrial dynamism in HepG2 KO cells which was more evident when both MBOAT7 and TM6SF2 genes were deleted. Then we performed a lentiviral transfection to overexpress *MBOAT7* and *TM6SF2* WT proteins alone or in combination in HepG2 KO cells. The WT overexpressed models strongly reduced the intracellular fat accumulation similarly to what was reported by Sharpe and colleagues in male C57BL6/J-MBOAT7^+/+^ mice and by Pant et al in Huh-7-TM6SF2 WT^+/+^ cells(39, 40). Interestingly, the WT overexpression of MBOAT7 or TM6SF2 in MBOAT7^-/-^TM6SF2^-/-^ clone brought out a glass effect resulting in a more conspicuous lipid reduction in MBOAT7^+/+^TM6SF2^-/-^ than MBOAT7^-/-^ TM6SF2^+/+^. This data confirmed the link between macro steatotic phenotype and MBOAT7 loss-of-function previously demonstrated in MBOAT7^-/-^cells, which resulted in the shift of phosphatidylinositols toward the synthesis of saturated and monounsaturated triacylglycerols with the consequent induction of DNL(30). Contrariwise, the *TM6SF2* KO induced the formation of smaller lipid droplets (LDs) by increasing the synthesis of triacylglycerols enriched in saturated and monounsaturated fatty acid chains(27).

As mentioned above, *TM6SF2* and *MBOAT7* KO models, more so when combined, impair mitochondrial morphology and biomass(27). Consistently, in the present study, we observed that the deletion of *MBOAT7* or *TM6SF2* alone and mainly together promoted fission while inhibited both fusion and mitophagy, showing an unbalanced mitobiogenesis and an increased assembly of misshapen and failed organelles as we recently demonstrated in hepatic biopsies of MASLD patients stratified according to genetic background (**doi:** https://doi.org/10.1101/2024.06.03.597155). Conversely, the *MBOAT7* and *TM6SF2* WT co-overexpression rebalanced the mitochondrial life course, thus ensuring a physiological organelles morphology, architecture, and activity. These results corroborate previous findings according to which the high number of mitochondria may correlate with impaired organelles’ dynamics and progressive MASLD(41–44). An increased mitochondrial biomass due to the accumulation of globular mitochondria in KO cell lines unlike the physiological *spaghetti-like* shaped in WT overexpressed cells, did not mirror a higher activity of the organelles. In keeping with this data, fibrotic mice fed with a high-trans-fat, high-fructose, and high-cholesterol (AMLN) diet, showed an increased number of disrupted mitochondria which were featured by low OXPHOS capacity, loss of cristae architecture and reduced expression of *Mnf1* and *Opa1*(45). In furtherance, Zhang et al demonstrated that *Mnf1* expression was reduced in 34 HCC patients and in MHCC97-H cells and inhibited cell proliferation, migration, and invasion, thereby playing a crucial role in impeding HCC development(46).

In another study, MASH patients showed high mitochondrial diameters, intra-mitochondria crystalline inclusions and granules in the matrix, which correlated with both mitochondrial swelling and OXPHOS failure(47). Previous evidence has reported that in hepatic tissue of patients and animal models with MASLD, the mitochondrial respiratory chain capacity is reduced. Compared to healthy individuals, patients with MASLD have a reduction in respiratory chain activity of 37% in complex I, 42% in complex II, 30% in complex III, 38% in complex IV, and 58% in complex V(48, 49). Consistently, we observed that *MBOAT7* and *TM6SF2* double KO decreased OXPHOS capacity, ATP production, ketogenesis and β-oxidation. The mitochondrial activity was rescued by the overexpression of the WT proteins and was paralleled by a reduced release of ccf-mtDNA. Moreover, KO cells promoted the tumorigenic switching in terms of anaerobic glycolysis, proliferation, and invasiveness, that was strongly attenuated in WT overexpressed models.

Notably, we found that the TM6SF2 loss-of-function more than MBOAT7 one may be mostly involved in the mitochondrial derangement by further increasing the total number of misshapen and failed mitochondria. In corroboration with our results, it has been demonstrated that the TM6SF2 deficiency reduced the amount of polyunsaturated fatty acids (PUFAs), along with alterations in mitochondrial β-oxidation in Huh-7 cells, whereas it induced changes in endoplasmic reticulum architecture in the small intestine of zebrafish, supporting that TM6SF2 loss-of-function impacts both organelles’ morphology and activity(50, 51). Concerning PNPLA3 polymorphism, Cas9 cells carrying I148M protein did not affect the mitochondrial dynamics. However, literature evidence reported that the overexpression of I148M in Huh-7 hepatoma cells was correlated with high levels of lactate and γ-glutamyl-amino acids, hallmarks of metabolic switching and mitochondrial dysfunction, respectively(36). Therefore, in order to investigate the possible crosslink between PNPLA3 loss-of-function and organelles’ maladaptation, we upregulated the I148M mutated protein in HepG2 cells in order to force the mutation effect and in Hep3B cells which were WT for the I148M variant (HepG2^I148M+/+^ and Hep3B^I148M+/+^).

The *in vitro* I148M overexpression enhanced LDs accumulation and reduced the mitochondrial functions in terms of OXPHOS rate and ATP reduction in both cell lines. This caused a boosted oxidative stress as well as release of lactate and ccf-mtDNA, suggesting the involvement of the I148M mutation in the mitochondrial failure.

To sum up, the dysfunctional metabolism encompassing lipid accumulation, oxidative stress and mitochondrial aberrances, alongside genetics lie at the center of MASLD pathogenesis(21). However, we firstly demonstrated a connection between MASLD-related genetics and mitochondrial dysfunction(52) and in this paper the *in vitro* results demonstrated that genetics impacts on mitobiogenesis thereby giving an explanation of higher organelles content and impaired activity.

High ccf-mtDNA levels in body fluids were correlated with advanced MASLD stages, underling their use as novel, non-invasive, cheaper, and repeatable MASLD diagnostic strategy(16). We observed in hepatoma cells that the silencing of MBOAT7 and TM6SF2 genes led to an enhanced release of ccf-mtDNA whose levels where conversely reduced in overexpressed cells. In addition, ccf-mtDNA correlated with the impaired respiration, in term of lower activity of complex V (ATP synthase) and reduced OCR.

Therefore, we tried to establish whether the circulating mitochondrial bioenergetic profile accurately reflects the hepatic one to support the potential use of molecule derived from damaged mitochondria (i.e. ccf-mtDNA) as biomarker of advanced disease in genetically predisposed individuals. Consistently with our *in vitro* findings, liver biopsies of patients carrying 3NRV exhibited a strong reduction of mitochondrial activity resulting in low OXPHOS rate. Notably, the PBMCs bioenergetic profile thoroughly reflected the hepatic one and showed a strong prognostic value for MASH-Fibrosis in 3NRV carriers (n=44)(22, 53, 54). Accordingly, serum respirometry was lower in 3NRV Fibroscan-MASLD patients, exhibiting again a sharp accuracy in predicting fibrosis. This evidence highlighted that serum OCR has a similar capacity to predict fibrosis in both biopsied and non-invasively assessed MASLD patients with a genetic predisposition. Additionally, the impaired serum respirometry seems to be specific of MASLD since its levels were higher in unrelated liver disease patients (n=45) and unchanged across different etiologies.

This study proves that restoring PNPLA3, MBOAT7, and TM6SF2 activities could counteract MASLD, pointing to them as potential therapeutic targets at the RNA level for personalized therapy. The targeting of PNPLA3 I148M variant in mice model by tri-antennary N-acetyl galactosamine (GalNAC3) conjugated with an antisense RNA oligonucleotide (ASO) decreased hepatic inflammation, steatosis, and fibrosis(55). In another recent study, targeting Pnpla3 in the 148M knock-in mice by Adeno associated viral AAV-mediated shRNA reduced hepatic triglyceride contents(56).

Focusing on our results, suitable approaches could be represented by gene therapy strategies based on the gene transfer that can introduce a WT copy to recover the function of the protein. For instance, the AAV-mediated delivery of a permanently active mutant form of human carnitine palmitoyl transferase 1A (hCPT1AM) in the liver of mice fed High Fat Diet enhanced liver fatty acid oxidation(57). Furthermore, the AAV8-mediated gene transfer approach that allows long-term hepatic SIRT1 overexpression counteracted high carbohydrate diet-induced MASLD and improved whole-body metabolism in adult mice(58). Supported by these literature evidence, we may delineate a new proof of concept regarding the applicability of gene transfer therapy in genetic-based MASLD management to prevent its progressive forms.

To conclude, the novel aspects of this study are twofold. We confirmed the contribute of *PNPLA3*, *MBOAT7*, and *TM6SF2* loss-of-function variants to mitochondrial dysfunction that features progressive MASLD, suggesting that their restoration which reverses the failure phenotype may be a successful therapeutic strategy. Secondly, we demonstrated for the first time that mitochondrial damage in PBMCs is the same as in the liver in MASLD patients and it is specific for the disease thus suggesting in a pioneering fashion the use of circulating mitochondrial biomarkers (i.e. mt-ccf, bioenergetic index) to foresee disease severity in genetically predisposed individuals.

## Fundings

This study was supported by Italian Ministry of Health (Ricerca Corrente 2024 - Fondazione IRCCS Cà Granda Ospedale Maggiore Policlinico), by Italian Ministry of Health (Ricerca Finalizzata Ministero della Salute GR-2019-12370172; RF-2021-12374481), PNRR-MCNT2-2023-12378295 and by 5x1000 2020 RC5100020B).

## Author Contributions

The authors’ responsibilities were as follows: EP, study design, data analysis and interpretation, manuscript drafting; ML and MM data analysis and interpretation; PP, MM, AQ data generation; ALF patients’ enrolment and manuscript revision; PD study design, manuscript drafting, data analysis and interpretation, funding acquisition, supervision and has primary responsibility for final content. All authors read and approved the final manuscript.

## Conflict of Interests

The authors declare that they have no conflict of interest.

## Abbreviations

Metabolic Dysfunction-Associated Steatotic Liver Disease (MASLD); metabolic dysfunction- associated steatohepatitis (MASH); peripheral blood mononuclear cells (PBMCs); mitochondria (mt); adenosine triphosphate (ATP); mitochondrial damage-associated molecular patterns (Mito-DAMPs); mitochondrial DNA (mtDNA) fragments (ccf-mtDNA); patatin-like phospholipase domain-containing 3 (PNPLA3);transmembrane 6 superfamily member 2 (TM6SF2); membrane bound o-acyltransferase domain-containing 7 (MBOAT7).

## Notes

### Competing Interest Statement

The authors have declared no competing interest.

### Summary of Updates

We have added experimental data. We have added another cohort of patients, in whom MASLD has been diagnosed non-invasively. Moreover, we have demonstrated the specificity of biomarkers for MASLD by assessing them in patients with other diseases.

