## Supplementary for "Circulating mitochondrial bioenergetics as fingerprint of the hepatic one: how to monitor genetic MASLD"

***Supplementary methods***

***Table S1:*** Clinical features of the MASLD biopsied patients (Discovery cohort)

| | **Discovery cohort (n=44)** |
|---|---|
| Sex, M | 29 (65.9) |
| Age, years | 52±13.89 |
| BMI, kg/m2 | 28.61±4.08 |
| IFG/T2D, yes | 16 (36.36) |
| HOMA-IR | 4.83±3.26 |
| Insulin, IU/ml | 22.27±17.80 |
| Total cholesterol, mmol/L | 5.01±0.98 |
| LDL cholesterol, mmol/L | 3.05±0.88 |
| HDL cholesterol, mmol/L | 1.31±0.28 |
| Triglycerides, mmol/L | 1.56±0.81 |
| ALT, IU/l | 4.04 {0.08-0.69} |
| AST, IU/l | 3.64{3.52-3.76} |
| Moderate/mild, NAS | 19 (43.2) |
| Severe, NAS | 25 (56.8) |
| **Number of Risk Variants (NRV)** | |
| 0 NRV | 10 (22.7) |
| GG PNPLA3 (1 NRV) | 11 (25.0) |
| TT MBOAT7 (1 NRV) | 9 (20.5) |
| TT TM6SF2 (1 NRV) | 7 (15.9) |
| 3 NRV | 7 (15.9) |

Values are reported as mean±SD, number (%) or median {IQR}, as appropriate. BMI: body mass index; IFG: impaired fasting glucose; T2D: type 2 diabetes; HDL: high density lipoprotein; LDL: low density lipoprotein; ALT: alanine aminotransferase; AST: aspartate aminotransferase; RV: risk variants. Characteristics of participants were compared across histological clinical phenotype using linear regression model (for continuous variables) or logistic regression model (for categorial characteristics). Variables with skewed distribution were logarithmically transformed before analyses. 0: indicates the absence of risk variants; 1NRV: indicates the presence of GG PNPLA3, TT MBOAT7 or CT TM6SF2 risk variants; 3 indicates the total number of risk variants carried.

***Table S2:*** **Clinical features of patients with non-invasive diagnosis of MASLD (*Fibroscan-MASLD cohort)***

| | **Fibroscan MASLD cohort (n=45)** |
|---|---|
| Sex, M | 31 (68.9) |
| Age, years | 50.76±11.04 |
| BMI, kg/m2 | 28.42±4.33 |
| IFG/T2D, yes | 3 (6.67) |
| HOMA-IR | 4.80±2-01 |
| Insulin, IU/ml | 17.14±8.45 |
| Total cholesterol, mmol/L | 2.27±011 |
| LDL cholesterol, mmol/L | 1.96±0.23 |
| HDL cholesterol, mmol/L | 1.68±0.13 |
| Triglycerides, mmol/L | 2.1±0.26 |
| ALT, IU/l | 1.64 {1.55-1.72} |
| AST, IU/l | 1.45{1.4-1.5} |
| CAP, db/m | 292±51.7 |
| Stiffness, kPa | 6.74±5.41 |
| **Number of Risk Variants (NRV)** | |
| 0 NRV | 9 (20.0) |
| GG PNPLA3 (1 NRV) | 10 (22.2) |
| TT MBOAT7 (1 NRV) | 9 (20.0) |
| CT/TT TM6SF2 (1 NRV) | 7 (15.6) |
| 3 NRV | 10 (22.22) |

Values are reported as mean±SD, number (%) or median {IQR}, as appropriate. BMI: body mass index; IFG: impaired fasting glucose; T2D: type 2 diabetes; HDL: high density lipoprotein; LDL: low density lipoprotein; ALT: alanine aminotransferase; AST: aspartate aminotransferase; CAP: controlled attenuation parameter; NRV: number of risk variants. Characteristics of participants were compared across histological clinical phenotype using linear regression model (for continuous variables) or logistic regression model (for categorial characteristics). Variables with skewed distribution were logarithmically transformed before analyses. 0 NRV: indicates the absence of risk variants; 1 NRV: indicates the presence of GG PNPLA3, TT MBOAT7 or CT/TT TM6SF2 risk variants; 3 NRV indicates the total number of risk variants carried.

***Table S3:*** **Clinical features of the patients belonging to the *Unrelated Liver Disease Cohort***

| | **Unrelated Liver Disease Cohort (n=44)** |
|---|---|
| Sex, M | 30 (68.18) |
| Age, years | 51±15.68 |
| BMI, kg/m2 | 24.96±4.51 |
| ALT, IU/l | 1.68 {1.45-1.90} |
| AST, IU/l | 1.63 {1.40-1.86} |
| CAP, db/m | 240.2±37.7 |
| Stiffness, kPa | 7.7±1.55 |

Values are reported as mean±SD, number (%) or median {IQR}, as appropriate. BMI: body mass index; ALT: alanine aminotransferase; AST: aspartate aminotransferase; CAP: controlled attenuation parameter. Characteristics of participants were compared across histological clinical phenotype using linear regression model (for continuous variables) or logistic regression model (for categorial characteristics).

***Lentiviral Overexpression***

MBOAT7 and TM6SF2 were overexpressed in MBOAT7-/- (MBOAT7+/+) and TM6SF2-/- (TM6SF2+/+) cells and then alone or in combination in MBOAT7-/-TM6SF2-/- clone (MBOAT7+/+TM6SF2-/-, MBOAT7-/-TM6SF2+/+, MBOAT7+/+TM6SF2+/+) through pLenti-C-mGFP-P2A-Puro lentiviral vectors, which were engineered to express a complete ORF fused with a GFP tag. We seeded 3∗104 cells in 24-well plates and they were incubated at 37ºC and 5% CO2 overnight. Multiplicity of infection was set at 2.5 and the number of lentiviral particles for the transduction were calculated according to the manufacturer's instructions (OriGene, Rockville, MD). Lentiviral particles were added to prewarmed cultured media for 24 hours. To introduce MBOAT7-GFP–tagged protein, MBOAT7-/- and MBOAT7-/-TM6SF2-/- cells were transduced with the LENG4 (MBOAT7) Human Tagged ORF Clone Lentiviral Particle (OriGene) containing a molecular sequence that aligns with the MBOAT7 mRNA (gene accession numbers: NM_024298.2 and NP_077274.2). To insert the TM6SF2-GFP, TM6SF2-/- and MBOAT7-/-TM6SF2-/- cells were transfected with TM6SF2 Human Tagged ORF Clone Lentiviral Particle (OriGene) carrying the molecular sequence that aligns with the TM6SF2 mRNA (gene accession numbers: NM_001001524.2 and NP_001001524.2). To embed MBOAT7-GFP and TM6SF2-GFP, MBOAT7-/-TM6SF2-/- cells were co-transduced with both LENG4 (MBOAT7) Human Tagged ORF Clone Lentiviral Particle (OriGene) and TM6SF2 Human Tagged ORF Clone Lentiviral Particle (OriGene). Stable cell lines carrying the WT forms of MBOAT7 and/or TM6SF2 (MBOAT7+/+, TM6SF2+/+, MBOAT7+/+TM6SF2-/-, MBOAT7-/-TM6SF2+/+, MBOAT7+/+TM6SF2+/+) were selected using the puromycin resistance gene (GE Healthcare Dharmacon, Inc).

***Oil Red O (ORO) staining***

Cells were plated on a 6-well plate (5 × 105 cells/well) in duplicate and left overnight in Dulbecco's modified Eagle medium (DMEM) containing 10% fetal bovine serum (FBS), 1% L-glutamine, and 1% penicillin/streptomycin. After 24 hours, growth media was removed and cells were kept for 24 hours in quiescent medium, containing 0.5% bovine serum albumin (BSA), 1% L-glutamine, and 1% penicillin/streptomycin. The day after, we performed ORO staining, which is a soluble red powder with high affinity for neutral triacylglycerols (TAGs) and lipids stored in the lipid droplets. Quiescent medium was removed, and the 6-well plates were gently rinsed with 2 mL sterile phosphate-buffered saline 1X. Next, cells were fixed with 4% formalin for 1 hour at room temperature. After fixation, each sample was washed with sterile water and 60% isopropanol was added for 5 minutes. Concurrently, we prepared ORO working solution by mixing 3 parts of ORO stock solution (300 mg of Red Oil powder in 100 mL di-isopropanol 100%) and 2 parts of sterile water, after filtration. ORO working solution (1 mL/well) was added to each sample and left for 40 minutes. Finally, plates were rinsed with tap water, paying attention not to disrupt the monolayer. LD content was visualized in pink-red color. The ORO-positive area was quantified by ImageJ software in 10 random micrographs (magnification, 200X) by calculating the ORO-positive area as a percentage of pixels above the threshold value with respect to the total pixels per area.

***Evaluation of Triglycerides and Cholesterol content***

A total of 2 x 106 of Cas9, MBOAT7-/-, TM6SF2-/-, MBOAT7-/- TM6SF2-/-, MBOAT7+/+, TM6SF2$^{+/+}$, MBOAT7$^{+/+}$TM6SF2$^{-/-}$, MBOAT7$^{-/-}$TM6SF2$^{+/+}$ and MBOAT7$^{+/+}$TM6SF2$^{+/+}$ cells were culture in DMEM containing 10% FBS, 1% L-glutamine, and 1% penicillin/streptomycin. After 24 hours, growth media was removed and cells were kept for 24 hours in quiescent medium, containing 0.5% bovine serum albumin (BSA), 1% L-glutamine, and 1% penicillin/streptomycin. The day after, cells were lysed in 0.5% Nonidet P-40 (NP-40) lysis buffer and cell lysates were used in quadruplicate to quantify the triglyceride and cholesterol content by exploiting the Triglycerides Colorimetric/Fluorometric Assay Kit and Cholesterol Colorimetric/Fluorometric Assay Kit, respectively. To measure triglyceride concentration, cells lysates were incubated with lipase for 20 minutes and then with the reaction mix for 30 minutes. The triglycerides will be converted to free fatty acids and glycerol. Glycerol is then oxidized to generate a product which reacts with a probe to generate color (spectrophotometry at $\lambda$= 570 nm) and fluorescence (Ex/Em = 535/587 nm). To assess cholesterol amount, cell lysates were incubated with the total cholesterol reaction mix with a specific probe for 60 minutes at 37 ºC generating color (570 nm) or fluorescence (Ex/Em = 538/587 nm).

***Gene Expression Analysis***

RNA was extracted from cell cultures using TRIzol reagent (Life Technologies–ThermoFisher, Carlsbad, CA). Total RNA (1 μg) was retrotranscribed with the VILO random hexamers synthesis system (Life Technologies–ThermoFisher, Carlsbad, CA). Quantitative real-time PCR was performed by an ABI 7500 fast thermocycler (Life Technologies), using the TaqMan Universal PCR Master Mix (Life Technologies–ThermoFisher, Carlsbad, CA) and TaqMan probes for human GAPDH, MBOAT7, TM6SF2, PGC1α, D-loop and Rnase P. SYBR Green chemistry (Fast SYBR Green Master Mix; Life Technologies–ThermoFisher, Carlsbad, CA) were exploited for the other genes. All reactions were delivered in quadruplicate. Data were normalized to β-actin and G3PDH housekeeping genes and results were expressed as mean and standard error (SE) and graphed as data points. Probes and primers are listed in **Tables S4A** and **S4B**, respectively.

**Table S4**. **A)** List of TaqMan Probes and **B)** sequence of primers used in Quantitative Real-Time PCR Experiments

**A)**

| PROBES | CATALOG NUMBER |
|---|---|
| **GAPDH** | THERMOFISHER HS02786621_G1 |
| **MBOAT** | THERMOFISHER HS00383302_M1 |
| **TM6SF2** | THERMOFISHER HS00403495_M1 |
| **PPARGC1α (PGC1α)** | THERMOFISHER #HS00173304_M1 |
| **MT-7S (D-LOOP)** | THERMOFISHER #HS02596861_S1 |
| **RNASE P** | THERMOFISHER #4401631 |

**B)**

| | FORWARD 5'→3' | REVERSE 5'→3' |
|---|---|---|
| ***B-ACTIN*** | GCTACAGCTTCACCACCACA | AAGGAAGGCTGGAAAAGAGC |
| ***MFN1*** | GAGGTGCTATCTCGGAGACAC | GCCAATCCCACTAGGGAGAAC |
| ***MFN2*** | CACATGGAGCGTTGTACCAG | TTGAGCACCTCCTTAGCAGAC |
| ***DRP1*** | ACCCGGAGACCTCTCATTCT | TGACAACGTTGGGTGAAAAA |
| ***FIS1*** | GATGACATCCGTAAAGGCATCG | AGAAGACGTAATCCCGCTGTT |
| ***PPARα*** | ATGGCATCCAGAACAAGGAG | TCCCGTCTTTGTTCATCACA |
| ***COXIII*** | TGACCCACCAATCACATGC | ATCACATGGCTAGGCCGGAG |
| ***ND1*** | CCCTAAAACCCGCCACATCT | GAGCGATGGTGAGAGCTAAGGT |
| ***DUSP1*** | CCTGACAGCGCGGAATCT | GATTTCCACCGGGCCAG |
| ***MCL1*** | CCAAGAAAGCTGCATCGAACCAT | CAGCACATTCCTGATGCCACCT |
| ***BAX*** | TTTGCTTCAGGGTTTCATCC | TCCTCTGCAGCTCCATGTTA |

***Western Blot Analysis***
Total protein lysates were extracted from cell cultures, using RIPA buffer containing 1 mmol/L Na-orthovanadate, 200 mmol/L phenylmethyl sulfonyl fluoride, and 0.02 µg/µL aprotinin. Samples were pooled prior to electrophoretic separation, and all reactions were performed in duplicate. Then, equal amounts of proteins (50 µg) were separated by SDS-PAGE, transferred electrophoretically to nitrocellulose membrane (BioRad, Hercules, CA, USA), and incubated with specific antibodies overnight. At least, three independent lots of freshly extracted proteins were used for experiments. The antibodies and concentration used are listed in **TableS5**.

**Table S5**. List of Antibodies and Relative Dilutions Used in Western Blot (WB) and Immunocytochemistry (ICC) Experiments

| ANTIBODY | CATALOG NUMBER |
|---|---|
| **Vinculin (1:1000 WB)** | ABCAM #AB73412 |
| **MBOAT7 (1:1000 WB)** | SIGMA-ALDRICH AV49811 |
| **TM6SF2 (1:1000 WB)** | THERMOFISHER PA5-69304 |
| **PGC1α (1:1000 WB; 1:200 ICC)** | NOVUS BIOLOGICALS NBP1-04676 |
| **MFN1 (1:1000 WB; 1:200 ICC)** | ABCAM #AB126575 |
| **MFN2 (1:200 ICC)** | ABCAM #AB56889 |
| **OPA1 (1:1000 WB; 1:200 ICC)** | ACAM #AB119685 |
| **DRP1 (1:1000 WB; 1:200 ICC)** | ABCAM #AB56788 |
| **PARKIN (1:1000 WB; 1:200 ICC)** | CELL SIGNALING #4211 |
| **PINK (1:1000 WB)** | CELL SIGNALING #6946 |
| **BNIP3-L (1:1000 WB)** | CELL SIGNALING #44060 |
| **BNIP3 (1:1000 WB)** | CELL SIGNALING #12396 |
| **LC3 (1:1000 WB)** | CELL SIGNALING #3868 |
| **PHOSPHO-UBIQUITIN (1:1000 WB)** | CELL SIGNALING #62802 |
| **OXPHOS (1:1000 WB)** | ABCAM #AB110411 |
| **COXI** | ABCAM AB110216 |
| **SDHA** | ABCAM AB110216 |

***Immunocytochemistry (ICC)***

A total of 1× $10^5$ cells were seeded on coverslips lodged in a 6-well plate in duplicate and kept overnight in DMEM containing 10% FBS, 1% L-glutamine, and 1% penicillin/streptomycin. After 24 hours, growth media was removed and cells were kept for 24 hours in quiescent medium, containing 0.5% bovine serum albumin (BSA), 1% L-glutamine, and 1% penicillin/streptomycin. Next, hepatocytes were fixed in 4% formalin for 10 minutes and permeabilized in 0.3% Triton X-100 (Sigma-Aldrich, St Louis, MO). Cells were incubated in 5% BSA for 30 minutes and with anti- primary antibody (**Table S5**) overnight at 4°C. Then, each sample was incubated with anti-rabbit/mouse horseradish-peroxidase–conjugated antibody, and 3,3′-diaminobenzidine was provided as chromogen. Nucleus were counterstained with hematoxylin. Finally, samples were mounted with a drop of aqueous VectaMount AQ Mounting Medium (Maravai LifeSciences, Inc, San Diego, CA).

***Mito Tracker staining***

A total of 1× $10^5$ cells were seeded on coverslips lodged in a 6-well plate in duplicate and kept overnight in DMEM containing 10% FBS, 1% L-glutamine, and 1% penicillin/streptomycin. After 24 hours, growth media was removed and cells were kept for 24 hours in quiescent medium, containing 0.5% bovine serum albumin (BSA), 1% L-glutamine, and 1% penicillin/streptomycin. The day after, cells were incubated with the MitoRed dye-working solution (Abcam, Cambridge, UK) for 30 minutes in a 37°C 5% CO2 incubator. Then, cells were washed twice with PBS 1X and fixed in 4% formalin for 20 minutes covered by lights. Finally, coverslips samples were mounted with a drop of aqueous VectaMount AQ Mounting Medium (Maravai LifeSciences, Inc, San Diego,

CA) and observed at confocal structured illumination microscopy (SIM) using a 100X oil immersion and pinhole of 1.2 to view the mitochondrial network number and morphology. The MitoRed-positive-area was quantified by ImageJ software in 10 random micrographs (magnification, 60X) by calculating the MitoRed-positive area as a percentage of pixels above the threshold value with respect to the total pixels per area. The MitoRed-positive-area of different mitochondrial shape was measured through ImageJ software in 10 random micrographs (magnification, 60X) by assessing 2 threshold value of MitoRed-positive area of *spaghetti*-like and globular mitochondria and then calculated the percentage of pixels above each threshold value with respect to the total pixels per area.

***Transmission Electron Microscopy (Tem)***
Cells were cultured as monolayer (70%–80% confluent) and trypsinized to obtain cell suspension. Cells were fixed with an aldehyde mixture (4% paraformaldehyde þ 2.5% glutaraldehyde in cacodylate buffer: pH 7.4) at 4C overnight. After primary fixation, cells were washed repeatedly with cacodylate buffer and were postfixed in 1% osmium tetroxide (OsO4) for 2 hours in the dark. Next, samples were left in 1.5% potassium ferrocyanide dissolved in 0.1 mol/L cacodylate for 1 hour on ice. Each sample was stained with 0.5% 5-uranyl acetate in water overnight. Finally, samples were dehydrated with increasing ethanol series, embedded in an Epon resin, and polymerized in an oven at 60C for 48 hours. Ultrathin (70–90 nm) sections were collected on nickel grids and observed with a Zeiss Leo 912 AB Omega TEM (Freiburg, Germany).

***Quantification of mtDNA (D-loop) content***
Genomic and mtDNA were extracted from cell lysates through QIAmp DNA Mini Kit (Manchester, UK). $2x10^6$ cells were resuspended in a Protease solution and incubated 56°C for 10 minutes to disruptprotein-DNAinteractions. Total DNA was trapped onto the QIAamp silica membrane, while contaminants were removed in the flowthrough. Subsequently, DNA was eluted in water and its concentration and quality were assessed by Nanodrop 1000 microvolume 42 spectrophotometer (ThermoFisher Scientific, U.S.A.). 5 ng/µl of DNA was used to quantify the amount of mtDNA. Specifically, we measured D-loop expression, the replication start site of the mtDNA, through TaqMan Assay. RNAse-P, a sequence known to exist in two copies in human genome, was used as a reference gene and both D-loop and RNase-P probes (**Table S4A**) run simultaneously. Median ΔCT per assay value was used as calibrator.

***Quantification of ccf-mtDNA content***
A total of 3 x $10^6$ of cells were culture in DMEM containing 10% FBS, 1% L-glutamine, and 1% penicillin/streptomycin. After 24 hours, growth media was removed and cells were kept for 24 hours in quiescent medium, containing 0.5% bovine serum albumin (BSA), 1% L-glutamine, and 1% penicillin/streptomycin. The day after, the medium was collected and concentrated with Vivaspin® 2 Centrifugal Concentrator (Sartorius, Göttingen, Germany) to extract and purified DNA fragments by using QIAmp DNA Blood Mini Kit (Manchester, UK). The Protease solution and AL Buffer were added to 150 µL concentrated cells media and incubated 56°C for 10 minutes to disrupt protein-DNA interactions. The DNA fragments up to 50 kb were trapped in columns providing silica membrane followed by two rinsing steps, while contaminants were removed in the flowthrough. Subsequently, DNA was eluted in water and its concentration and quality were assessed by Nanodrop 1000 microvolume 42 spectrophotometer (ThermoFisher Scientific, U.S.A.). 20 ng/µl of DNA was used to quantify the amount of ccf-mtDNA. Quantitative real-time PCR was performed by an ABI 7500 fast thermocycler (Life Technologies), using SYBR Green chemistry (Fast SYBR Green Master Mix; Life Technologies–ThermoFisher, Carlsbad, CA) and COXIII and ND1 primers listed in **Table S4B**. Specifically, the thermocycling setup was set as follow: 95°C for 10 minutes, 40 cycle: 95°C for 10 minutes, 60°C for 1 minute, 95°C for 15 second, 60°C for 1 minute and 95°C for 15 second. All reactions were delivered in quadruplicate. Data were normalized on the standard curve obtained from serial dilutions of a sample pool at known concentration to measure the quantity (gram) of DNA. Then, the amount of ccf-mtDNA (gram/150 µL), encompassing the sum of COXIII and ND1 quantities, was divided with the size of PCR fragments (bp; 102 bp COXIII, 109 bp ND1) and the molar mass per base pair ($g\ mol^{-1}$)[59]. The product was finally multiplied with Avogadro's constant to obtain the quantity (picogram) of DNA fragments expressed as logarithmic. Results were represented as mean and standard error (SE) and graphed as data points.

***Evaluation of Mitochondrial Complex I activity***
A total of 5mg/ml of NAFLD patients liver biopsies and PBMCs homogenate in MAS buffer was loaded in a 96-well plate in quadruplicate. The mitochondrial complex I enzyme activity was measured through a colorimetric assay at 450 nm at 20 second-1min intervals for up to 30 minutes with shake between readings at room temperature by using an Assay solution (Abcam, Cambridge, UK).

***Evaluation of Mitochondrial Complex III activity***

A total of 3 x $10^6$ of cells were culture in DMEM containing 10% FBS, 1% L-glutamine, and 1% penicillin/streptomycin. After 24 hours, growth media was removed and cells were kept for 24 hours in quiescent medium, containing 0.5% bovine serum albumin (BSA), 1% L-glutamine, and 1% penicillin/streptomycin. The day after, mitochondria were isolated from cell lysates by exploiting a Mitochondria Isolation Kit for Cultured Cells (Abcam, Cambridge, UK). The mitochondria were resuspended in RIPA buffer and 1.5 to 10 μg was seeded in a 96-well plate in quadruplicate. Additionally, 15 μg of NAFLD patients liver biopsies and PBMCs homogenate in MAS buffer were loaded in a 96-well plate in quadruplicate. The mitochondrial complex III activity was measured through a colorimetric assay at 550 nm at 30 second intervals for up to 10 minutes at room temperature by using a Sample Mix alone or in combination with Antimycin A, specific inhibitor of the mitochondrial complex III (Abcam, Cambridge, UK).

***Evaluation of Mitochondrial Complex IV activity***
Mitobiogenesis In-Cell ELISA Kit Abcam (Cambridge, UK) exploits a quantitative immunocytochemistry (ICC) to measure subunit I of mitochondrial COXI (Complex IV) and nuclear DNA (nDNA)-encoded succinate dehydrogenase complex flavoprotein subunit A (SDHA) protein levels in live cells. Briefly, cells are seeded in a 96-well plate ($3x10^5$/well) in quadruplicate, fixed with 4% paraformaldehyde and permeabilized through 1X Triton-X 100. Targets of interest were detected with highly specific, well-characterized cocktail of monoclonal antibodies, which were incubated overnight at 4°C. Then, AP-labelled and HRP-labeled secondary antibodies were used to generate a colorimetric reaction that could be measured at 405 and 600 nm, respectively.

***Evaluation of Mitochondrial Complex V activity***
A total of 3 x $10^6$ of cells were culture in DMEM containing 10% FBS, 1% L-glutamine, and 1% penicillin/streptomycin. After 24 hours, growth media was removed and cells were kept for 24 hours in quiescent medium, containing 0.5% bovine serum albumin (BSA), 1% L-glutamine, and 1% penicillin/streptomycin. The day after, mitochondria were isolated from cell lysates by exploiting a Mitochondria Isolation Kit for Cultured Cells (Abcam, Cambridge, UK) and resuspended in RIPA buffer. The ATP Synthase Enzyme Activity Microplate Assay Kit (Abcam, Cambridge, UK) exploits a quantitative ELISA assay to measure the mitochondrial Complex V by using Capture antibodies that are pre-coated in the wells of premium Nunc MaxiSorp™ modular microplates. 10 μg/50 μL of extracted mitochondria and 15 μg of NAFLD patients liver biopsies and PBMCs homogenate in MAS buffer were added in quadruplicate on the plate and incubated for 3 hours at room temperature. Then, empty the wells and incubate the Lipid Mix for 45 minutes at room temperature. Finally, the Reagent Mix was added to generate a colorimetric reaction that could be measured at 340 nm at 30°C for 60-120 minutes.

***Evaluation of Citrate Synthase activity***
A total of 3 x 106 of cells were culture in DMEM containing 10% FBS, 1% L-glutamine, and 1% penicillin/streptomycin. After 24 hours, growth media was removed and cells were kept for 24 hours in quiescent medium, containing 0.5% bovine serum albumin (BSA), 1% L-glutamine, and 1% penicillin/streptomycin. The day after, mitochondria were isolated from cell lysates by exploiting a Mitochondria Isolation Kit for Cultured Cells (Abcam, Cambridge, UK). The mitochondria were resuspended in RIPA buffer and 1-50 μL were added in a 96 well plate as well as 5 μl of NAFLD patients liver biopsies and PBMCs homogenate in MAS buffer in quadruplicate for the Citrate Synthase Assay Kit (Abcam, Cambridge, UK). The Reaction Mix was used to measure absorbance at 412 nm at 25°C for 20-40 minutes.

***Evaluation of Lactate Dehydrogenase enzymatic activity***
A total of 3 x $10^6$ of cells were culture in DMEM containing 10% FBS, 1% L-glutamine, and 1% penicillin/streptomycin. After 24 hours, growth media was removed and cells were kept for 24 hours in quiescent medium, containing 0.5% bovine serum albumin (BSA), 1% L-glutamine, and 1% penicillin/streptomycin. The day after, it was performed the Lactate Dehydrogenase (LDH) Assay Kit (Abcam, Cambridge, UK) to measure the LDH enzymatic activity. Cells were homogenate with Assay Buffer in ice, centrifuge at 4°C at 10,000 x g for 15 minutes and collect the supernatant. 50 μL of cell lysates in quadruplicate were added in 96 well and incubated with a Reaction Mix containing the LDH assay buffer and LDH substrate mix to measure the absorbance at 450 nm for 30-60 minutes at 37°C.

***Evaluation of Lactate Dehydrogenase production***
A total of 3 x $10^6$ of cells were culture in DMEM containing 10% FBS, 1% L-glutamine, and 1% penicillin/streptomycin. After 24 hours, growth media was removed and cells were kept for 24 hours in quiescent medium, containing 0.5% bovine serum albumin (BSA), 1% L-glutamine, and 1% penicillin/streptomycin. The day after, the medium was collected and concentrated with Vivaspin® 2 Centrifugal Concentrator (Sartorius, Göttingen, Germany) to measure the LDH activity by Lactate Dehydrogenase (LDH) Assay Kit (Abcam, Cambridge, UK) which exploits the conversion of lactate into pyruvate and NADH. 50 μL of concentrated culture

media was plated in quadruplicate in 96 well and then, the reaction mix containing the LDH assay buffer, the PicoProbe and LDH substrate mix was added to measure the fluorescence reaction at Ex/Em 535/587 nm for 10-30 minutes at 37°C.

***Mito Stress Seahorse Assay***

The Agilent Seahorse XF Cell Mito Stress Test measures parameters of mitochondrial function by directly quantifying the oxygen consumption rate (OCR) of cells on the Seahorse XFe and XF Extracellular Flux Analyzers. The assay uses the built-in injection ports on XF sensor cartridges to add modulators of respiration into cell well during the test that are Oligomycin, Carbonyl cyanide-4 (trifluoromethoxy) phenylhydrazone (FCCP), Rotenone, and Antimycin. $5x10^4$ cells in quadruplicate were plated on a Seahorse XF24-well microplate in DMEM containing 10% FBS, 1% L-glutamine, and 1% penicillin/streptomycin. After 24 hours, the growth media was replaced by the assay medium, supplemented with glucose, pyruvate, and glutamine, and cells were incubated at 37°C for 60 minutes. Then, Seahorse XF24-well microplate was adapted on the pre-warmed XF sensor cartridges congaing the Oligomycin, FCCP, Rotenone and Antimycin drugs and the OCR was measured through XFe24 analyzer, with 3 baseline measurements recorded before and after adding several drugs (1.5 µM Oligomycin, 0.5 µM FCCP and 0.5 µM Rotenone + Antimycin).

***Glycolytic Seahorse Assay***

The Agilent Seahorse XF Glycolysis Stress Test is the standard assay for measuring glycolytic function in cells by directly measuring the extracellular acidification rate (ECAR) and assesses the key parameters of glycolytic flux: Glycolysis, Glycolytic Capacity, Glycolytic Reserve, as well as non-glycolytic acidification. The assay uses the built-in injection ports on XF sensor cartridges to add modulators of glycolysis into cell well during the test that are Oligomycin and 2-deoxy-glucose (2-DG), a glucose analog. $5x10^4$ cells in quadruplicate were plated on a Seahorse XF24-well microplate in DMEM containing 10% FBS, 1% L-glutamine, and 1% penicillin/streptomycin. After 24 hours, the growth media was replaced by the glycolysis stress test medium without glucose or pyruvate and cells were incubated at 37°C for 60 minutes. Then, Seahorse XF24-well microplate was adapted on the pre-warmed XF sensor cartridges congaing oligomycin and 2-DG to measure ECAR through XFe24 analyzer, with 3 baseline measurements recorded before and after adding several drugs (5 µM Oligomycin and 500 mM 2-DG).

***Evaluation of ROS/RNS production***

The Cellular ROS/RNS Detection Assay Kit (Abcam, Cambridge, UK) is designed to directly monitor real time reactive oxygen and/or nitrogen species (ROS/RNS) production in live cells using fluorescence microscopy. Cells are seeded in a 96-well plate ($3x10^5$/well) in quadruplicate in DMEM containing 10% FBS, 1% L-glutamine, and 1% penicillin/streptomycin to ensure 50- 70% confluency on the day of the experiment. Then, growth media was removed, and cells were kept for 24 hours in quiescent medium, containing 0.5% bovine serum albumin (BSA), 1% L-glutamine, and 1% penicillin/streptomycin. The day after, cells and 10 µg of NAFLD patients liver biopsies and PBMCs homogenate in MAS buffer were loaded with the ROS/RNS 3-Plex Detection Mix containing Oxidative Stress Detection Reagent, Superoxide Detection Reagent and NO Detection Reagent for 2 hours. For the last 30 minutes, cells were treated with the NO Inducer (L-Arginine) and the ROS Inducer (Pyocyanin). After rinsing the cells, they could be observed under fluorescent/confocal microscope or it could detect oxidative stress (ROS) and superoxide (RNS) at Ex/Em 490/525nm and 550/620nm, respectively.

***Evaluation of apurinic/apyrimidinic (AP) sites***

A total of $3 \times 10^6$ of cells were culture in DMEM containing 10% FBS, 1% L-glutamine, and 1% penicillin/streptomycin. After 24 hours, growth media was removed and cells were kept for 24 hours in quiescent medium, containing 0.5% bovine serum albumin (BSA), 1% L-glutamine, and 1% penicillin/streptomycin. The day after, genomic DNA was extracted from cell lysates through QIAmp DNA Mini Kit (Manchester, UK). Cells lysates were resuspended in a Protease solution and incubated 56°C for 10 minutes to disruptprotein-DNAinteractions. Total DNA was trapped onto the QIAamp silica membrane, while contaminants were removed in the flowthrough. Subsequently, DNA was eluted in water and its concentration and quality were assessed by Nanodrop 1000 microvolume 42 spectrophotometer (ThermoFisher Scientific, U.S.A.). 100 µg/mL of purified genomic DNA was used for the DNA damage – AP sites – Assay Kit (Abcam, Cambridge, UK) to detect aldehyde site (AP sites) at 450 nm.

***Evaluation of Malondialdehyde production***

A total of $3 \times 10^6$ of cells were culture in DMEM containing 10% FBS, 1% L-glutamine, and 1% penicillin/streptomycin. After 24 hours, growth media was removed and cells were kept for 24 hours in quiescent medium, containing 0.5% bovine serum albumin (BSA), 1% L-glutamine, and 1% penicillin/streptomycin. The day after, cells were lysed in 0.5% Nonidet P-40 (NP-40) lysis buffer and cell lysates were used in quadruplicate to perform the Lipid Peroxidation (MDA) Assay Kit (Abcam, Cambridge, UK). Cell lysates were incubated at room temperature for 30-60 minutes with the MDA color reagent solution and then measure the absorbance at 695-700 nm.

***Wound Healing Assay***
Cells were plated on 6-well plate ($8 \times 10^5$/well) and incubated with DMEM containing 10% FBS, 1% L-glutamine, and 1% penicillin/streptomycin. A fine scratch was introduced using a sterile pipette tip in a monolayer of cells at approximately 90% confluency. The wounds were photographed (objective, 100×) at 24 and 48 hours. Each experiment was performed in triplicate.

***Cell Proliferation Assay***
$2.5 \times 10^2$ cells/well were seeded in a 96-well plate in quadruplicate and incubated in DMEM containing 10% FBS, 1% L-glutamine, and 1% penicillin/streptomycin overnight. Cell proliferation was measured at baseline using the CellTiter96-Aqueous One Solution Cell Proliferation Assay (MTS:3-(4,5-dimethylthiazol-2-yl)-5-(3-carboxymethoxyphenyl)-2-(4-sulfophenyl)-2H-tetrazolium)) kit (Promega Corporation, Fitchburg, WI). Fresh growth media was provided for 24-48-72 h and 1 week. MTS reagent (20 µL/well) was added to the cells followed by incubation for 4 hours in a 5% CO2 humidified incubator at 37°C, and the absorbance was measured at 490 nm, at 0, 24, 48, and 74 hours, and after 1 week. At least 3 independent experiments were performed.

***Invasion Assay***
The invasion assay[17] exploits the transwell insert membrane coated with Collagen (3 mg/mL) above which $25 \times 10^3$ cells were plated in duplicate in DMEM containing 10% FBS, 1% L-glutamine, and 1% penicillin/streptomycin. The underside of the transwell was filled with culture media. After 24 hours, growth media was removed and cells were kept for 24 hours in quiescent medium, containing 0.5% bovine serum albumin (BSA), 1% L-glutamine, and 1% penicillin/streptomycin. After 24 and 48 hours, cells were fixed in 4% formalin for 10 minutes. The transwell membrane containing the fixed cells was cut and mounted with a drop of aqueous VectaMount AQ Mounting Medium on a cover slide. Cells were photographed (objective, 100×) at 24 and 48 hours. Each experiment was performed in triplicate.

***Supplementary results***

***The lentiviral overexpression of MBOAT7 and/or TM6SF2 wild-type genes in CRISPR-Cas9 knock-out models***

MBOAT7 and TM6SF2 deficiency in HepG2 cells led to mitochondrial aberrances. Therefore, to explore whether their recovery could have an impact on mitochondrial number, morphology and function we leveraged a lentiviral transfection in $MBOAT7^{-/-}$, $TM6SF2^{-/-}$ and $MBOAT7^{-/-}TM6SF2^{-/-}$ cells by using pLenti-C-mGFP-P2A-Puro lentiviral vectors containing the puromycin selection aiming to obtain stable cell lines ($MBOAT7^{+/+}$, $TM6SF2^{+/+}$, $MBOAT7^{+/+}TM6SF2^{-/-}$, $MBOAT7^{-/-}TM6SF2^{+/+}$, $MBOAT7^{+/+}TM6SF2^{+/+}$). The vector deigns embraces a GFP tag fused with an ORF targeting the protein which should be upregulated (OriGene Technologies, Inc., PS100093). The messenger RNA (mRNA) and protein levels of MBOAT7 and TM6SF2 significantly increased in $MBOAT7^{-/-}$ (**Figure S1A-B**) and $TM6SF2^{-/-}$ (**Figure S1C-D**) cells alone or in combination in $MBOAT7^{-/-}TM6SF2^{-/-}$ ones (**Figure S1E-F**), thus restoring the CRISPR-Cas9-driven deletion which mimics the human protein loss-of-functions (adjusted $^{**}p<0.01$ *vs* $MBOAT7^{-/-}$, $TM6SF2^{-/-}$, $MBOAT7^{-/-}TM6SF2^{-/-}$; adjusted $^{*}p<0.05$ *vs* $MBOAT7^{-/-}$).
Since MBOAT7 and TM6SF2 natural functional role allows to lipid assembly, metabolism, and transport, we evaluated the intracellular LDs in KO and overexpressed models by exploiting the ORO staining. Lipid accumulation was significantly lower in $MBOAT7^{+/+}$ and $TM6SF2^{+/+}$ cells compared to $MBOAT7^{-/-}$ and $TM6SF2^{-/-}$, respectively, showing a similar Cas9 phenotype. Despite both MBOA $T7^{+/+}TM6SF2^{-/-}$ and $MBOAT7^{-/-}TM6SF2^{+/+}$ models exhibited a strong lipid reduction than $MBOAT7^{-/-}TM6SF2^{-/-}$ clone, deletion of MBOAT7 or TM6SF2 alone depicts a mirror effect on LDs size. Indeed, MBOAT7 KO in $MBOAT7^{-/-}TM6SF2^{+/+}$ clone led to larger LDs size due to their enrichment in saturated/monounsaturated TAG [30]. Otherwise, TM6SF2 KO in $MBOAT7^{+/+}TM6SF2^{-/-}$ cell line promotes the assembly of small LDs by overexpressing saturated diacylglycerols (DAGs) [27]. The greatest reduction in intracellular fat was detected in $MBOAT7^{+/+}TM6SF2^{+/+}$ model, thus suggesting that the restore of both MBOAT7 and TM6SF2 proteins rather than one could protect against steatosis (**Figure S1G**). According to the qualitative results, the measurement of ORO-positive area (**Figure S1H**) alongside intracellular triglycerides (**Figure S1I**) and cholesterol (**Figure S1J**) content significantly decreased in all the overexpressed cells compared to the KO ones (adjusted $^{**}p<0.01$ *vs* Cas9, $MBOAT7^{-/-}$, $TM6SF2^{-/-}$, $MBOAT7^{-/-}TM6SF2^{-/-}$).

***Proliferation and invasiveness drop after the overexpression of MBOAT7 and/or TM6SF2 wild-type genes***

To investigate the involvement of MBOAT7 and TM6SF2 loss-of-functions in the malignant transformation and to deepen how their WT overexpression could attenuate the carcinogenesis, we examined cells proliferation and invasiveness. The scratch test revealed that at 48 hours the wound healing was improved in $MBOAT7^{-/-}$, $TM6SF2^{-/-}$ and completely patched up in $MBOAT7^{-/-}TM6SF2^{-/-}$cell lines. Conversely, all overexpressed cells showed a slight proliferative phenotype (**Figure S4A**). In keeping with these results, the MTS assay (**Figure S4B**) highlighted the increased proliferation rate from 48 hours peaking at 1 week in $MBOAT7^{-/-}$, $TM6SF2^{-/-}$ and $MBOAT7^{-/-}TM6SF2^{-/-}$ models compared to their overexpressed ones, exhibiting the higher growth rate in $MBOAT7^{-/-}TM6SF2^{-/-}$ clone detectable at 48 and even more at 72 hours (adjusted *$p<0.05$, **$p<0.01$ *vs* Cas9, $MBOAT7^{-/-}$, $TM6SF2^{-/-}$, $MBOAT7^{-/-}TM6SF2^{-/-}$). Inversely, the WT upregulation of *MBOAT7* and *TM6SF2* alone and in combination significantly reduced the growth capacity from 24 hours up to 1 week. According to these data, mRNA levels of *Dual Specificity Phosphatase 1* (*DUSP1*) (**Figure S4C**) and *Induced Myeloid Leukemia Cell Differentiation Protein* (*MCL1*) (**Figure S4D**), genes acting in proliferation and anti-apoptotic processes, respectively, augmented in $MBOAT7^{-/-}$, $TM6SF2^{-/-}$ and $MBOAT7^{-/-}TM6SF2^{-/-}$ cell lines whereas decreased after *MBOAT7* and/or *TM6SF2* WT overexpression, (adjusted *$p<0.05$ vs $MBOAT7^{-/-}TM6SF2^{-/-}$; **$p<0.01$ *vs* Cas9, $MBOAT7^{-/-}$, $TM6SF2^{-/-}$, $MBOAT7^{-/-}TM6SF2^{-/-}$). Otherwise, the expression of *Bcl-2-associated X protein* (*BAX*) (**Figure S4E**), a pro-apoptotic regulator, was downregulated after the deletion of *MBOAT7* and/or *TM6SF2* genes while strongly rose beyond their WT overexpression (**$p<0.01$ *vs* $MBOAT7^{-/-}$, $TM6SF2^{-/-}$, $MBOAT7^{-/-}TM6SF2^{-/-}$). Finally, we investigated cell migration by exploiting cells aptitude to pass across transwell insert membrane covered by collagen (**Figure S4F**). At 48 hours the number of migrated cells increased in $MBOAT7^{-/-}$ cells and even more in $TM6SF2^{-/-}$ and $MBOAT7^{-/-}TM6SF2^{-/-}$ models showing high cells quantity already at 24 hours. In comparison, the WT overexpression of *MBOAT7* and *TM6SF2* attenuated the invasiveness at both 24 and 48 hours displaying the hugely effect in $MBOAT7^{+/+}TM6SF2^{+/+}$ clone (**$p<0.01$ *vs* $MBOAT7^{-/-}$, $TM6SF2^{-/-}$, $MBOAT7^{-/-}TM6SF2^{-/-}$). Therefore, the restore of MBOAT7 and TM6SF2 WT functions in KO models could ameliorate the tumorigenic phenotype by reducing the proliferation capacity and invasiveness.

***Table S6. Association between NRV=3 and hepatic oxidative stress, citrate synthase activity, mitochondrial complexes activity in the Discovery cohort.***

| **Liver biopsies** | **ROS/RNS** | | **$H_2O_2$** | |
|---|---|---|---|---|
| | **β {95%CI}** | **P value** | **β {95%CI}** | **P value** |
| Age, Years | -3.30{-12.08-5.47} | 0.45 | -0.12{-0.35-0.11} | 0.30 |
| Sex, M | -136.80 {-258.15- -15.45} | **0.027** | -4.38 {-7.59- -1.17} | 0.08 |
| BMI, Kg/m2 | 4.99 {-23.71-33.70} | 0.72 | 0.42 {-0.34-1.17} | 0.27 |
| IFG/T2D, yes | 47.21 {-79.25-173.68} | 0.45 | -0.24 {-3.58-3.10} | 0.88 |
| **NRV=3** | 267.58 {110.84-4242.31} | **0.0011** | 4.07 {-0.06-8.22} | **0.05** |
| | **Citrate synthase** | | **Mitochondrial complex I** | |
| Age, Years | 0.0003 {-0.0002-0.0009} | 0.26 | 0.0014 {0.0002-0.002} | **0.015** |
| Sex, M | 0.001 {-0.006- -0.009} | 0.72 | -0.012 {-0.028-0.003} | 0.11 |
| BMI, Kg/m2 | 0.0013 {-0.0005-0.003} | 0.15 | 0.003 {-0.0003-0.0037 | 0.07 |
| IFG/T2D, yes | -0.007 {-0.15- -0.0008} | 0.07 | 0.002 {-0.014-0.018} | 0.77 |
| **NRV=3** | -0.01 {0.0018- -0.02} | **0.02** | -0.037 {-0.058- -0.017} | **0.0004** |
| | **Mitochondrial complex III** | | **Mitochondrial complex V** | |
| Age, Years | -0.0006{-0.0004-0.0005} | 0.08 | -0.0008{-0.0015-0.0001} | **0.017** |
| Sex, M | 0.004 {-0.002-0.01} | 0.17 | -0.003 {-0.12-0.0005} | 0.47 |
| BMI, Kg/m2 | 0.0003 {-0.0012-0.002} | 0.62 | 0.0013 {-0.0007-0.003} | 0.20 |
| IFG/T2D, yes | 0.005 {-0.001-0.013} | 0.10 | 0.013 {0.004-0.022} | **0.005** |
| **NRV=3** | -0.011 {-0.02- -0.003} | **0.009** | -0.046 {-0.058- -0.035} | **<0.0001** |

The co-presence of the PNPLA3, MBOAT7 and TM6SF2 variants (NRV=3) was associated with hepatic ROS/RNS production (P =0.0011, β = 267.58, 95 % CI, 110.84 to 4242.31), $H_2O_2$ release (P=0.05, β =4.07, 95%CI, -0.06 to 8.22), low citrate synthase activity (P<0.0001, β = -0.02, 95%CI, -0.03 to -0.01), low mitochondrial complex I activity (P=0.0004, β = -0.037, 95%CI, -0.058 to -0.017); low mitochondrial complex III activity (P=0.009, β = -0.011, 95%CI, -0.02 to -0.003) and low ATP synthase (mitochondrial complex V) activity (P <0.0001, β = -0.046, 95 % CI, -0.058 to -0.035) in the Discovery cohort at generalized linear model adjusted for age, sex, body mass index (BMI) and impaired fasting glucose/type 2 diabetes (IFG/T2D), Values are reported as mean±SD, number (%) or median {IQR}, as appropriate. BMI: body mass index; IFG/T2D: impaired fasted glucose/ type 2 diabetes; NRV: number of risk variants; β: beta coefficients; CI: confidence interval. p<0.05 was considered statistically significant.

***Table S7. Association between NRV=3 and PBMCs oxidative stress, citrate synthase activity, mitochondrial complexes activity in the Discovery cohort.***

| PBMCs | ROS/RNS | | $H_2O_2$ | |
|---|---|---|---|---|
| | **β {95%CI}** | **P value** | **β {95%CI}** | **P value** |
| Age, Years | -2.97{-12.63-6.69} | 0.54 | 0.018 {-0.19-0.22} | 0.85 |
| Sex, M | 47.44 {-77.04-171.92} | 0.44 | 0.56 {-2.15-3.28} | 0.67 |
| BMI, Kg/m2 | 49.86 {20.80-78.91} | **0.0011** | 1.11 {0.47-1.74} | **0.0009** |
| IFG/T2D, yes | -31.61 {-162.63-99.40} | 0.63 | -1.52 {-4.38-1.33} | 0.28 |
| NRV=3 | 1.85 {-157.29-161.01} | 0.98 | 0.50 {-2.98-3.96} | 0.77 |
| | **Citrate synthase** | | **Mitochondrial complex I** | |
| Age, Years | 0.0009 {0.0003-0.001} | **0.001** | 0.0006 {-0.0007-0.002} | 0.36 |
| Sex, M | -0.004 {-0.011-0.003} | 0.25 | -0.0019 {-0.019-0.016} | 0.82 |
| BMI, Kg/m2 | 0.0011 {-0.0006-0.002} | 0.22 | 0.0052 {0.001-0.009} | **0.013** |
| IFG/T2D, yes | -0.0038 {-0.011-0.004} | 0.33 | -0.011 {-0.03-0.007} | 0.24 |
| **NRV=3** | -0.02 {-0.03- -0.01} | **<0.0001** | -0.02 {-0.048- -0.003} | **0.02** |
| | **Mitochondrial complex III** | | **Mitochondrial complex V** | |
| Age, Years | -0.0006 {-0.0004-0.0003} | 0.74 | 0.0003{-0.0002-0.0009} | 0.0003 |
| Sex, M | -0.007 {-0.12- -0.001} | **0.009** | 0.001 {-0.006-0.009} | 0.72 |
| BMI, Kg/m2 | 0.0003 {-0.0016-0.0008} | 0.52 | 0.0013 {-0.0005-0.003} | 0.15 |
| IFG/T2D, yes | 0.001 {-0.004-0.006} | 0.68 | -0.007 {-0.015-0.0008} | 0.07 |
| **NRV=3** | -0.02 {-0.03- -0.01} | **<0.0001** | -0.01 {0.0018-0.022} | **0.02** |

The co-presence of the PNPLA3, MBOAT7 and TM6SF2 variants (NRV=3) in the PBMCs of the Discovery cohort was associated with ROS/RNS and $H_2O_2$ production, low citrate synthase activity (P=0.02, β = -0.01, 95%CI, 0.0018 to -0.02), low mitochondrial complex I activity (P=0.02, β = -0.02, 95%CI, 0.048 to -0.003), low mitochondrial complex III activity (P<0.0001, β = -0.02, 95%CI, -0.03 to -0.01), low ATP synthase (mitochondrial complex V) activity (P=0.02, β = -0.01, 95%CI, 0.0018 to -0.022) at generalized linear model adjusted for age, sex, body mass index (BMI) and impaired fasting glucose/type 2 diabetes (IFG/T2D),. Values are reported as mean±SD, number (%) or median {IQR}, as appropriate. BMI: body mass index; IFG/T2D: impaired fasted glucose/ type 2 diabetes; NRV: number of risk variants; β: beta coefficients; CI: confidence interval. p<0.05 was considered statistically significant.

***Table S8. Association between 3NRV and reduced respiration (complex I/IV) both in liver and PBMCs in the Discovery cohort***

| **Seahorse complex I/IV** | **Liver biopsies** | | **PBMCs** | |
|---|---|---|---|---|
| | **β {95%CI}** | **P value** | **β {95%CI}** | **P value** |
| Age, Years | 0.68 {-0.49-1.86} | 0.25 | 0.78 {-0.35-1.92} | 0.17 |
| Sex, M | -19.37 {-36.10- -2.65} | **0.02** | -28.54 {-43.75- -13.33} | **0.0004** |
| BMI, Kg/m2 | 3.81 {0.15-7.47} | 0.051 | 2.95 {-0.26-6.17} | 0.07 |
| IFG/T2D, yes | 5.29 {-12.13-22.72} | 0.54 | -12.92 {-28.95-3.09} | 0.11 |
| **NRV=3** | -43.37 {-64.81- -21.92} | **0.0001** | -30.7 {-49.96- -11.45} | **0.002** |

Values are reported as mean±SD, number (%) or median {IQR}, as appropriate. BMI: body mass index; IFG/T2D: impaired fasted glucose/ type 2 diabetes; NRV: number of risk variants; β: beta coefficients; CI: confidence interval. p<0.05 was considered statistically significant. Generalized linear model adjusted for age, sex, BMI, and IGF/T2D and NRV=3.

***Table S9. Association between 3NRV and reduced respiration (complex II/IV) both in liver and PBMCs in the Discovery cohort***

| **Seahorse complex II/IV** | **Liver biopsies** | | **PBMCs** | |
|---|---|---|---|---|
| | **β {95%CI}** | **P value** | **β {95%CI}** | **P value** |
| Age, Years | -0.051 {-1.23-0.21} | 0.16 | -0.50 {-0.78- -0.22} | 0.57 |
| Sex, M | -0.16 [60] | 0.97 | 1.72 {-2.27-5.71} | 0.39 |
| BMI, Kg/m2 | 0.42 {-1.83-2.67} | 0.70 | 0.098 {-0.77-0.97} | 0.82 |
| IFG/T2D, yes | -4.14 {-14.89-6.59} | 0.44 | -2.27 {-6.45-1.89} | 0.27 |
| **NRV=3** | -27.24 {-10.47- -14.02} | **0.0001** | -20.08 {-25.20- -14.96} | **<.0001** |

Values are reported as mean±SD, number (%) or median {IQR}, as appropriate. BMI: body mass index; IFG/T2D: impaired fasted glucose/ type 2 diabetes; NRV: number of risk variants; β: beta coefficients; CI: confidence interval. p<0.05 was considered statistically significant. Generalized linear model adjusted for age, sex, BMI, and IGF/T2D and NRV=3.

***Table S10. Association between 3NRV and reduced PBMCs respiration (complex I/IV) in the Fibroscan-MASLD cohort***

| **Seahorse complex I/IV** | **PBMCs Fibroscan-MASLD cohort** | |
|---|---|---|
| | **β {95%CI}** | **P value** |
| Age, Years | 0.27 {-1.16-1.71} | 0.70 |
| Sex, M | 19.73 {-13.01-52.48} | **0.23** |
| BMI, Kg/m2 | -1.85 {-5.38-1.67} | 0.29 |
| IFG/T2D, yes | -18.96 {-78.3-40.36} | 0.52 |
| **NRV=3** | -63.06 {-85.42- -40.7} | **<.0001** |

Values are reported as mean±SD, number (%) or median {IQR}, as appropriate. BMI: body mass index; IFG/T2D: impaired fasted glucose/ type 2 diabetes; NRV: number of risk variants; β: beta coefficients; CI: confidence interval. p<0.05 was considered statistically significant. Generalized linear model adjusted for age, sex, BMI, and IGF/T2D and NRV=3.

***Table S11. Association between 3NRV and reduced PBMCs respiration (complex I/IV) in the overall MASLD cohort***

Values are reported as mean±SD, number (%) or median {IQR}, as appropriate. BMI: body mass index; IFG/T2D: impaired fasted glucose/ type 2 diabetes; NRV: number of risk variants; β: beta coefficients; CI: confidence interval. p<0.05 was considered statistically significant. Generalized linear model adjusted for age, sex, BMI, and IGF/T2D and NRV=3.

| **Seahorse complex I/IV** | **PBMCs Overall MASLD cohort** | |
|---|---|---|
| | **β {95%CI}** | **P value** |
| Age, Years | 0.37 {-0.70-1.45} | 0.50 |
| Sex, M | 22.05 {-3.39-47.50} | 0.08 |
| BMI, Kg/m2 | -0.07 {-2.63-2.48} | 0.95 |
| IFG/T2D, yes | 9.23 {-18.71-37.18} | 0.50 |
| **NRV=3** | -79.05 {-96.29- -61.82} | **<.0001** |
| Age, Years | 0.59 {-0.55-1.74} | 0.30 |
| Sex, M | 17.90 {-9.93-45.74} | 0.20 |
| BMI, Kg/m2 | 0.26 {-2.44-2.98} | 0.84 |
| IFG/T2D, yes | 3.51 {-26.22-33.24} | 0.81 |
| **PNPLA3 G alelle, yes** | -7.35 {-20.94- -6.22} | 0.28 |
| **MBOAT7 T alelle, yes** | -30.46 {-44.64- -16.28} | **<.0001** |
| **TM6SF2 T alelle, yes** | -29.02 {-43.76- -14.28} | **0.0002** |

***Table S12. Specific association between reduced serum OCR and MASLD***

| Seahorse complex I/IV | β {95%CI} | P value |
|---|---|---|
| Age, Years | 0.27 {-1.16-1.71} | 0.70 |
| Sex, M | 19.73 {-13.01-52.48} | 0.23 |
| **Biopsied-MASLD *vs* Unrelated MASLD** | -63.06 {-85.42- -40.7} | **<.0001** |
| Age, Years | -0.05 {-0.96-0.85} | 0.89 |
| Sex, M | 14.1 {-11.05-39.25} | 0.26 |
| **Fibroscan-MASLD *vs* Unrelated MASLD** | -20.29 {-31.82- -8.75} | **0.0008** |
| Age, Years | -0.39 {-0.71-1.50} | 0.47 |
| Sex, M | 9.59 {-20.50-39.68} | 0.52 |
| **Overall MASLD *vs* Unrelated MASLD** | -16.61 {-31.76--1.46} | **0.03** |

Values are reported as mean±SD, number (%) or median {IQR}, as appropriate. β: beta coefficients; CI: confidence interval. $p<0.05$ was considered statistically significant. Generalized linear model adjusted for age, sex.

***Supplementary legend figures***

**Figure S1.** ***MBOAT7 and/or TM6SF2 WT overexpression improves lipid overload due to their silencing.*** **A-C-E)** The expression of *MBOAT7* and *TM6SF2* was evaluated by reverse-transcription quantitative PCR and normalized to the β-actin housekeeping gene. **B-D-F)** Protein levels of MBOAT7 and TM6SF2 tagged with GFP (MBOAT7-GFP and TM6SF2-GFP) were assessed by Western blot and normalized to the vinculin housekeeping gene. **G**) LDs accumulation was assessed by ORO staining (magnification, 630×). **H)** ORO positive (+ve) areas were quantified by ImageJ in 6 random nonoverlapping micrographs per condition by calculating the percentage of pixels above the threshold value in respect to total pixels per area. **I)** The intracellular triglycerides amount was measured by exploiting the Triglycerides Colorimetric/Fluorometric Assay. **J)** Total cholesterol was measured in cell lysates by using the Cholesterol Colorimetric/Fluorometric Assay Kit. At least 3 independent experiments were conducted. For bar graphs, data are expressed as means and SE. Adjusted $*P < .05$ and $**P < .01$.

***Figure S2. MBOAT7 and/or TM6SF2 WT overexpression rebalances the mitochondrial lifecycle and turnover.*** **A-E*)*** The mRNA levels of *PGC1*a, *Mfn1*, *Mfn2*, *Drp1* and *Fis1* was evaluated by reverse-transcription quantitative PCR and normalized to the β-actin housekeeping gene. **F-I)** Cytoplasmatic and nuclear localization of Mfn1, Mfn2, Opa1 and Drp1 were represented by Immunocytochemistry pictures. At least 3 independent experiments were conducted. For bar graphs, data are expressed as means and SE. Adjusted $*P < .05$ and $**P < .01$

***Figure S3. MBOAT7 and/or TM6SF2 WT overexpression in knock-out models restore the mitochondrial functions.*** **A)** The complex III enzymatic activity was measured through a colorimetric assay in isolated mitochondria. **B)** the MTCO1 (mtDNA– encoded subunit of the complex IV) activity was measured by Mitobiogenesis In-Cell ELISA Kit in isolated mitochondria from cell lysates. **C)** The ATP5A (mtDNA–encoded subunit of the complex V) activity was quantified by ATP Synthase Enzyme Activity Microplate Assay Kit in isolated mitochondria from cell lysates. **D)** The citrate synthase activity was assessed through the Citrate Synthase Assay Kit in isolated mitochondria from cell lysates. At least 3 independent experiments were conducted. For bar graphs, data are expressed as means and SE. Adjusted $*P < .05$ and $**P < .01$.

**Fig S4.** ***MBOAT7 and/or TM6SF2 WT restore in knock-out cells attenuates the tumorigenic phenotype.*** **A)** Representative images of wound healing assay were acquired at 0, 24 and 48 hours (magnification, 100×). The dotted lines indicate the scratch width. **B)** The proliferation rate was examined through MTS assay for 0, 24, 48,

72 hours and 1 week (λ = 490 nm). **C-D-E)** The expression of *DUSP1, MCL1* and *BAX* was evaluated by reverse-transcription quantitative PCR and normalized to the β-actin housekeeping gene. **F)** Representative images of invasiveness assay were acquired at 24 and 48 hours (magnification, 100×). The total number of cells were quantified by ImageJ in 10 random nonoverlapping micrographs per condition. At least 3 independent experiments were conducted. For bar graphs, data are expressed as means and SE. Adjusted $*P < .05$ and $**P < .01$.

### Figure S1

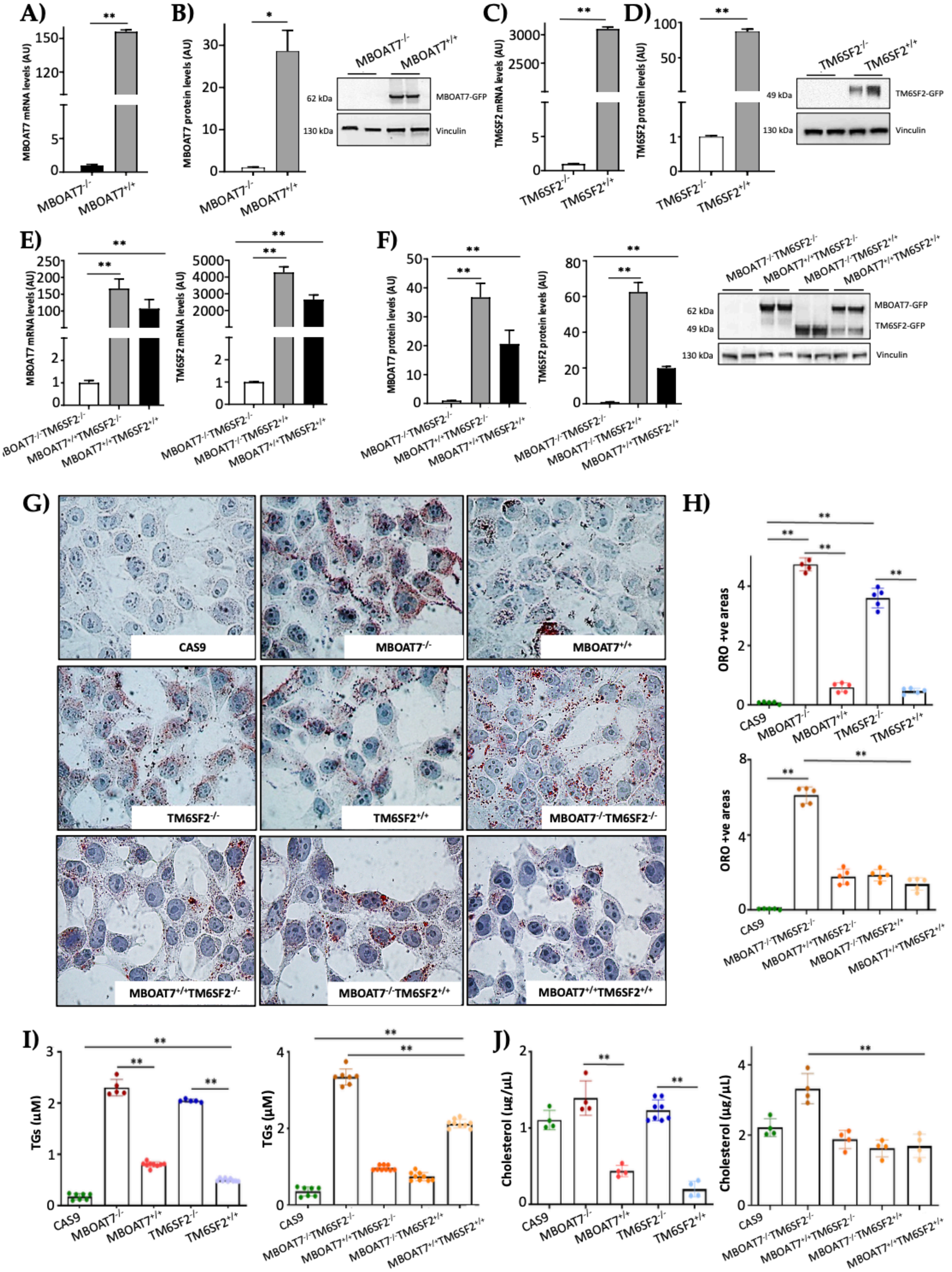

A)
B)
C)
D)
E)
F)
G)
H)
I)
J)
MBOAT7 mRNA levels (AU)
MBOAT7 protein levels (AU)
TM6SF2 mRNA levels (AU)
TM6SF2 protein levels (AU)
MBOAT7-GFP
TM6SF2-GFP
Vinculin
62 kDa
49 kDa
130 kDa
CAS9
MBOAT7-/-
MBOAT7+/+
TM6SF2-/-
TM6SF2+/+
MBOAT7-/-TM6SF2-/-
MBOAT7+/+TM6SF2-/-
MBOAT7-/-TM6SF2+/+
MBOAT7+/+TM6SF2+/+
ORO +ve areas
TGs (μM)
Cholesterol (μg/μL)

### Figure S2

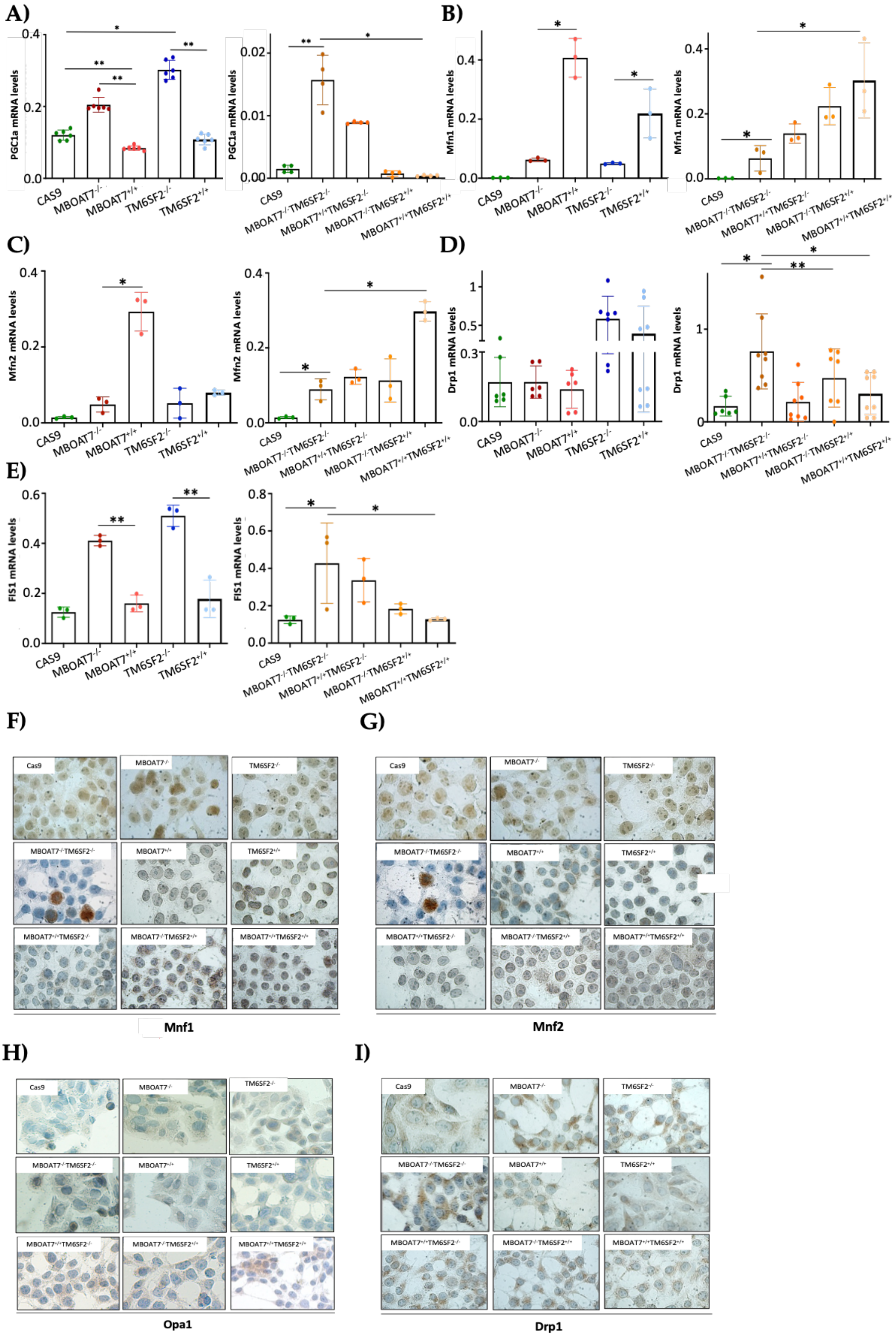
A)
B)
C)
D)
E)
F)
G)
H)
I)
PGC1a mRNA levels
Mfn1 mRNA levels
Mfn2 mRNA levels
Drp1 mRNA levels
FIS1 mRNA levels
CAS9
MBOAT7-/-
MBOAT7+/+
TM6SF2-/-
TM6SF2+/+
MBOAT7-/-TM6SF2-/-
MBOAT7+/+TM6SF2-/-
MBOAT7-/-TM6SF2+/+
MBOAT7+/+TM6SF2+/+
Cas9
Mnf1
Mnf2
Opa1
Drp1

### Figure S3

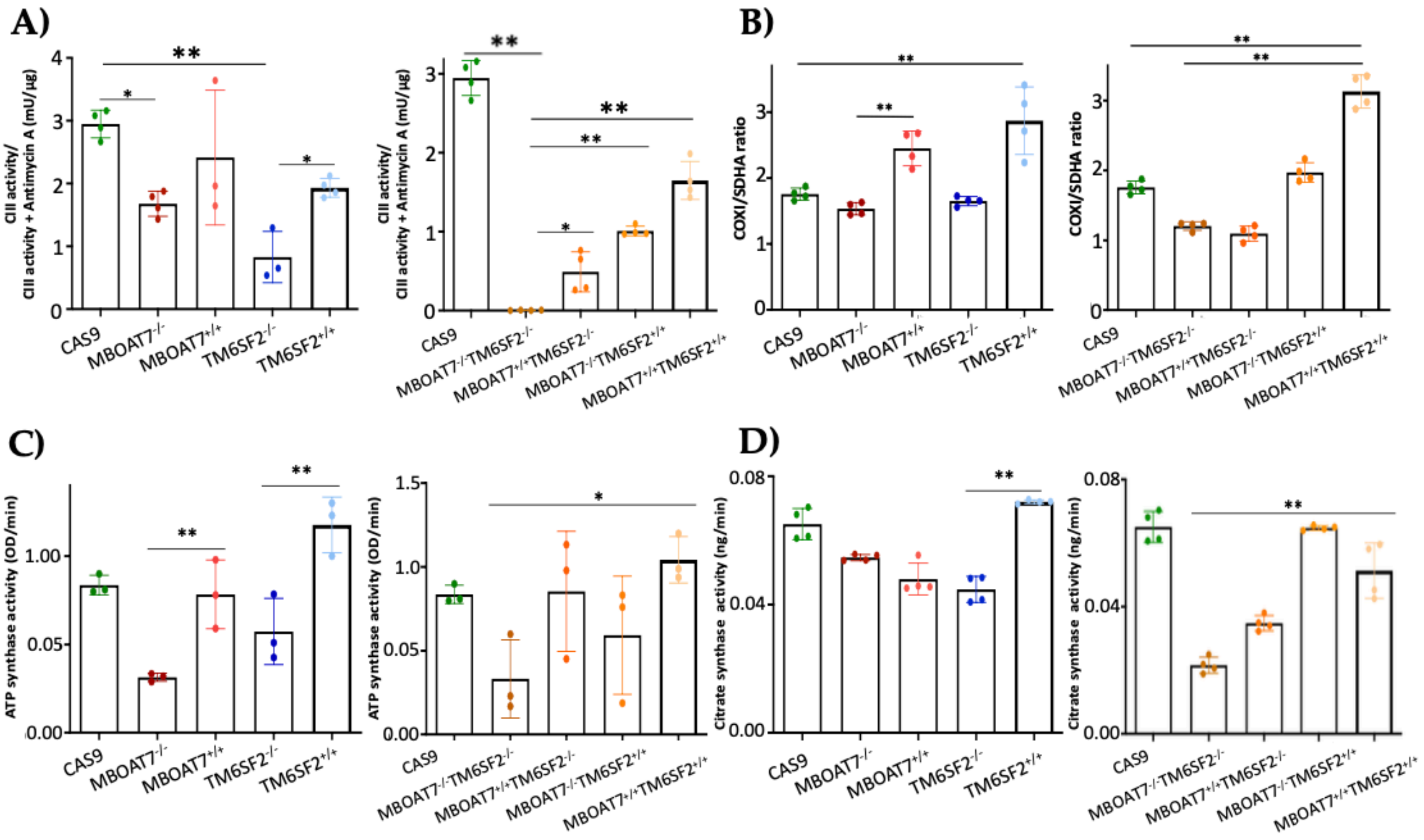

A)
B)
C)
D)
CII activity/ CII activity + Antimycin A (mU/µg)
COXI/SDHA ratio
ATP synthase activity (OD/min)
Citrate synthase activity (ng/min)
CAS9
MBOAT7-/-
MBOAT7+/+
TM6SF2-/-
TM6SF2+/+
MBOAT7-/-TM6SF2-/-
MBOAT7+/+TM6SF2-/-
MBOAT7-/-TM6SF2+/+
MBOAT7+/+TM6SF2+/+

### Figure S4

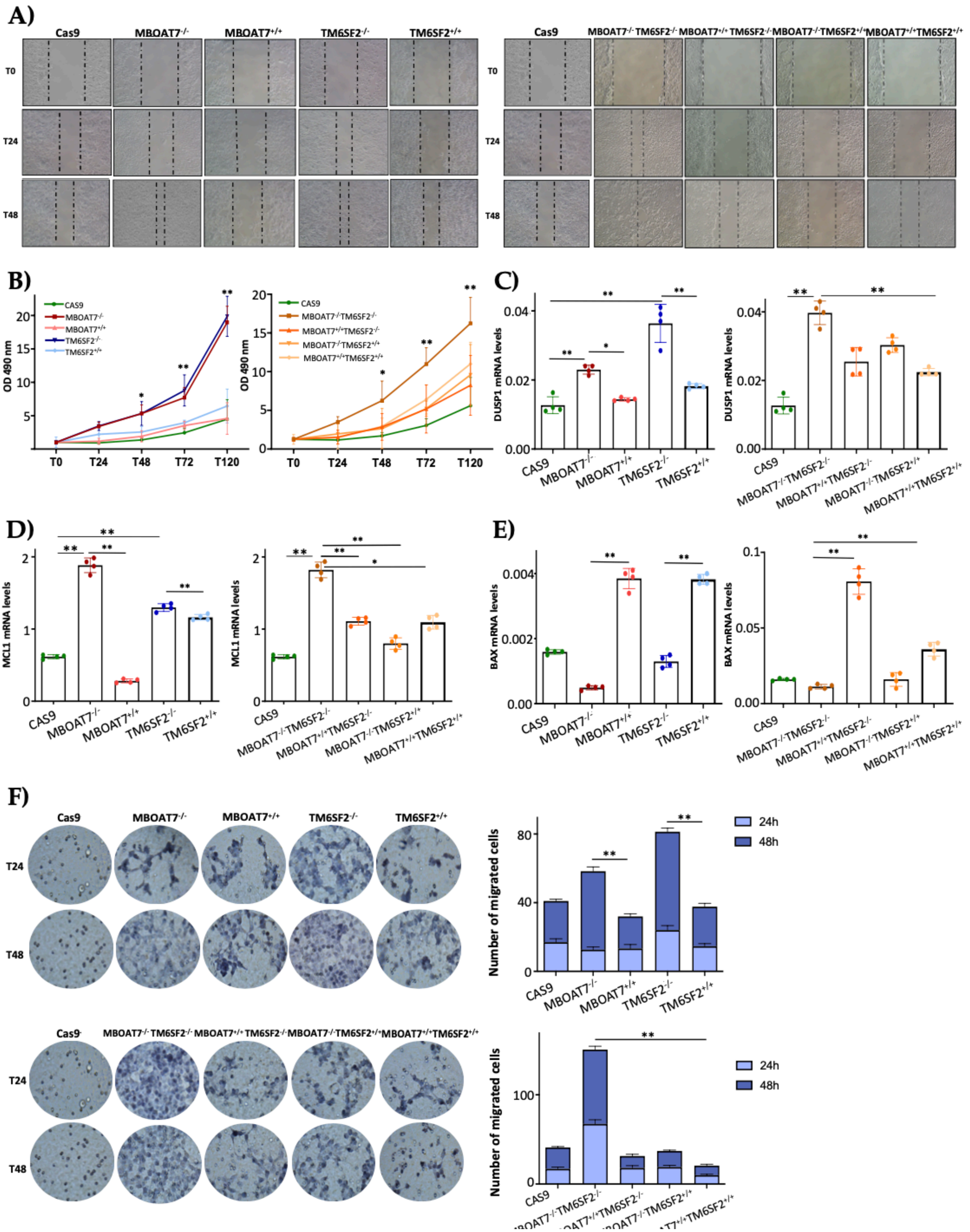
A)
Cas9
MBOAT7-/-
MBOAT7+/+
TM6SF2-/-
TM6SF2+/+
MBOAT7-/-TM6SF2-/-
MBOAT7+/+TM6SF2-/-
MBOAT7-/-TM6SF2+/+
MBOAT7+/+TM6SF2+/+
T0
T24
T48
B)
OD 490 nm
CAS9
T72
T120
C)
DUSP1 mRNA levels
D)
MCL1 mRNA levels
E)
BAX mRNA levels
F)
Number of migrated cells
24h
48h
